# Spatio-temporal evaluation of drought adaptation in wheat revealed NDVI and MTSI as powerful tools for selecting tolerant genotypes

**DOI:** 10.1101/2023.01.29.526148

**Authors:** S. Srinatha Reddy, G. Mahendra Singh, Uttam Kumar, Pradeep Bhati, Manish Vishwakarma, Sudhir Navathe, K. J. Yashavanthakumar, Ramesh Chand, Sandeep Sharma, Vinod K. Mishra, Arun K. Joshi

## Abstract

**Context:** Water stress is fast becoming a major limiting factor for wheat production. Hence, identifying drought tolerant genotypes is critical for sustaining the food supply chain. However, there are no phenotypic markers or statistical models available that may be employed for the efficient selection of field grown drought tolerant wheat genotypes.

**Objectives:** We evaluated wheat genotypes to (1) identify novel sources of drought tolerance (2) understand underlying adaptation mechanisms of drought tolerance (3) identify phenotypic markers and a stable model for the selection of drought tolerant genotypes

**Methods:** One hundred ninety-six diverse wheat genotypes were evaluated at three different locations in India: Banaras Hindu University (BHU), Varanasi (E1 and E3: control; E2 and E4: drought); Agharkar Research Institute (ARI), Pune (E5 and E7: control; E6 and E8: drought) and Borlaug Institute for South Asia (BISA), Jabalpur (E9 and E11: control; E10 and E12: drought) for various agronomic, physiological and yield traits for two consecutive years. Drought was imposed at the heading stage (Z59) by withholding irrigation for four weeks until the moisture reading reached <45% than the control (100%).

**Results:** The performance of all genotypes significantly declined under drought at all the locations. Normalized difference vegetation index (NDVI) significantly correlated (r = 0.41** and 0.36**) with the grain yield under drought during maturity. At the same time, there was no association under control conditions (r = 0.07 and 0.10) at the BHU center during 2020-21 and 2021-22, respectively. Stress indices, such as geometric mean productivity (GMP) and stress tolerance index (STI), showed a high correlation (r= 0.89** and r = 0.88**, respectively) with the grain yield under drought and were effective in differentiating drought tolerant genotypes. GGE bi-plots discriminated the environments (observed obtuse angle between E3 with E6 and E9, E4 with E6 and E9) having negative relation and cross-over interaction for grain yield. Further, the multi-trait stability index (MTSI) identified 29 stable genotypes across all environments and was predicted as the most accurate model due to its fewer Root Mean Square Prediction Difference (RMSPD) values.

**Conclusion:** NDVI is a useful high throughput screening tool under drought and MTSI is an effective method for selecting stable wheat genotypes across different water stressed locations.

**Implications:** The identified tools (NDVI), method (MTSI), and tolerant genotypes appear to be valuable resources that together will be useful in the ongoing breeding programs to enhance the drought tolerance of wheat.

**Highlights:**

- Drought stress significantly reduces wheat production globally.
- NDVI serves as a useful tool for high throughput screening under drought for field-based experiments.
- AMMI, GGE, WAASBY and MTSI plots are effective in detecting GEI effects.
- MTSI is a reliable tool for selecting stable and high mean performers.

## 1. Introduction

Wheat is one of the major staple cereals grown worldwide [1]. It is the most cultivated crop globally in terms of area (219 million hectares, 2020) [2]. Global wheat production in 2020 accounted for 760 million tonnes and stood as the second most-produced cereal after maize [2]. Current climate change patterns are predicted to increase drought and flood prone areas, which could significantly threaten global food security [3].

Wheat has been experiencing drought stress in most cultivating areas [4]. The periodic drought affects more than 50% of cultivated areas [5,6]. Drought being multidimensional stress profoundly impacts the physiological and morphological process and ultimately affects grain yield [7]. Flowering and grain filling are considered critical stages, and the incidence of drought (terminal drought) at this stage will profoundly impact yield [8]. Since it interferes with pollen sterility, grain number and size affect the wheat quality and productivity [9]. Wheat crops reported varying levels of drought-tolerant mechanisms. It varied with their growth stages [10], resulting in changes in cell wall integrity, photosynthetic rate, stomatal closure, and higher levels of oxidative stress [10]. Physiological traits serve yield stability parameters in evaluating wheat genotypes under drought conditions [11]. Genotypes with higher flag leaf photosynthesis during the grain filling period are likely to produce better grain yield than those with lower flag leaf photosynthesis [12]. Thus, understanding changes in morpho-physiological traits associated with drought is important to select and develop better drought adopting genotypes [13]. However, such phenotypic markers are limited in use for screening tolerance under water stress.

Drought tolerance is classified as a complex trait; its complexity is further intensified with more significant environmental influence [14]. India has diverse agro-climatic regions, influenced by a wide range of climatic patterns causing uneven and unpredictable levels of drought stress [15]. Identifying drought-tolerant genotypes that perform stably across environments is an essential consideration in the present scenario [16]. In dealing with current and future climatic changes, it is necessary to exploit genetic diversity to breed drought-tolerant wheat genotypes. Genetic diversity is usually a major source of novel alleles for stress adaptations and understanding “adaptation mechanisms” could enhance our insights into plant response to environmental stresses [17]. A few studies have been done that utilize genetic diversity to enhance the drought tolerance in wheat, [18,19]; however, a detailed understanding of drought adaptation response in wheat is still lacking.

Over the years, with the advancement in statistical tools, several models have been put forward to understand genotypic responses under various environmental stress [20]. Drought tolerant indices have been useful to classify drought tolerant genotypes [21]. Identifying drought-tolerant genotypes and testing across different environments helps in determining the genotypes true genetic potential [22]. Multi-trait stability index (MTSI) provides a unique and more reliable method of selection, where it considers multiple traits to obtain a higher mean performer and stable genotype [23,24]. MTSI has been utilized in selecting drought and salinity tolerant genotypes in soybean [25] and drought tolerant genotypes in chickpea [26]. However, the identification of drought tolerant genotypes using MTSI has not been reported in wheat.

This study was conducted to evaluate and identify stable lines from a diverse panel of wheat using different indices and models with the following objectives: (i) To identify drought tolerant lines across different locations and understand their adaptation mechanism, (ii) To employ MTSI as a unique selection strategy that relies on, both, mixed and fixed effects.

## 2. Materials and methods

### 2.1 The initial screening and planting material

The present study used a diverse panel of 294 spring wheat genotypes (Wheat Associated Mapping Initiative; WAMI Panel) representing 28 countries [27]. Initial screening was done to select genotypes with similar heading dates (one week interval) to avoid biases in plant response against drought due to differences in developmental stages. A total of 196 genotypes out of 294 were found to have similar heading dates and were chosen for the experiment.

### 2.2 Multi-environmental evaluation

One hundred ninety-six selected wheat genotypes from the WAMI panel were evaluated in the two consecutive years *Rabi* 2020-21 and 2021-22 at three locations, i.e., Institute of Agricultural Sciences (IAS), BHU, Varanasi; Agharkar Research Institute (ARI), Pune, Maharashtra; Borlaug Institute of South Asia (BISA), Jabalpur, Madhya Pradesh. Details of the test environments are given in Supplementary Table 1. Geographical information, including coordinates and altitude of three test locations, is shown in Fig. 1a.

**Fig. 1.**
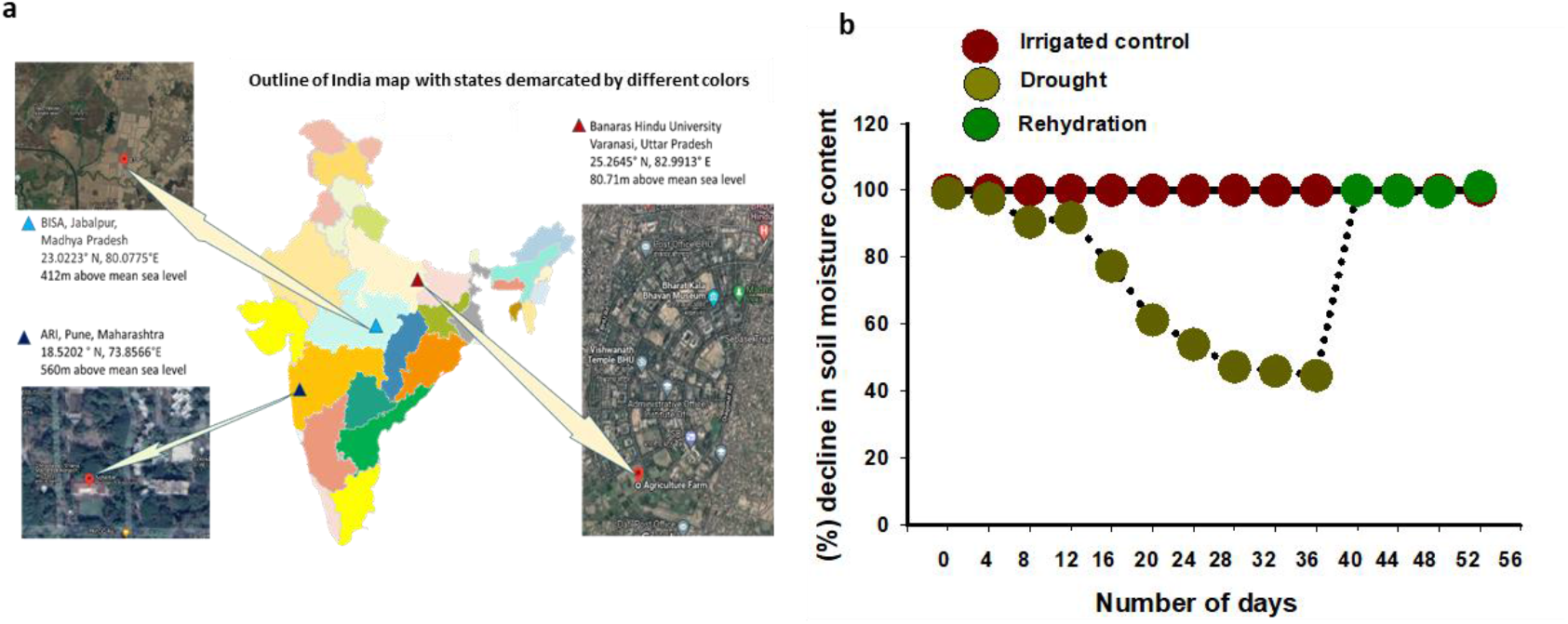
(a). Geographical information of testing environments for 196 WAMI genotypes at BHU, Varanasi; ARI, Pune and BISA, Jabalpur. (b) Percent decline in soil moisture content in a time series of drought treatment as compared to irrigated control at BHU center during *Rabi* 2021-22.

Trials were conducted in alpha lattice design with two replications. Each genotype was sown in 1 m row with 20 cm spacing between the rows. To induce drought stress, irrigation was stopped at the heading stage (Z59) [28], and the treatment was prolonged for four weeks until the soil moisture recorded 40-45% under drought treated plots than control (100%) at all the locations. Soil moisture observations were recorded every fourth day during drought treatment (Fig. 1b) using a soil moisture meter (LUTRON Soil Moisture Meter PMS-714, Lutron electronic enterprise co., ltd., Taiwan). The irrigation was resumed after the treatment period and followed regular agronomic practices till the harvest of the crop.

Traits such as the number of days to heading (DH), days to maturity (DM), plant height (PH), thousand kernel weight (TKW), and grain yield (GY) were recorded. Normalized Difference Vegetative Index (NDVI) was recorded using a handheld Green seeker, Trimble agriculture, during heading (Z59), grain filling (Z77), and maturity (Z91) stages. Leaf chlorophyll content was recorded using SPAD (Soil Plant Area Development, SPAD-502Plus, Konica Minolta, Inc., Japan) at heading, grain filling and maturity stages. Canopy temperature (CT) was also recorded using a handheld infrared Thermometer (LT300, Sixth sense) at all the three stages. CTD (Canopy Temperature Depression) was estimated after measuring canopy temperature and by following the formula [29,30].

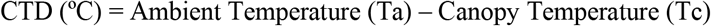

### 2.3 Stress indices

Drought tolerance was estimated for grain yield data by calculating the stress indices. The stress indices and correlation table were analyzed in Microsoft Excel 2016. Stress indices calculated are as follows:

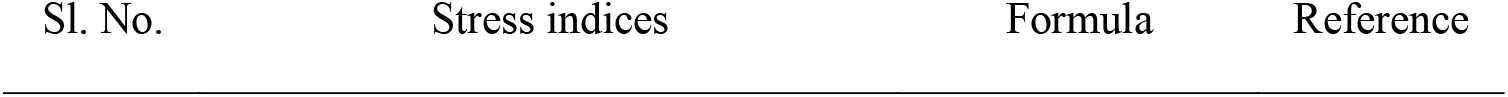
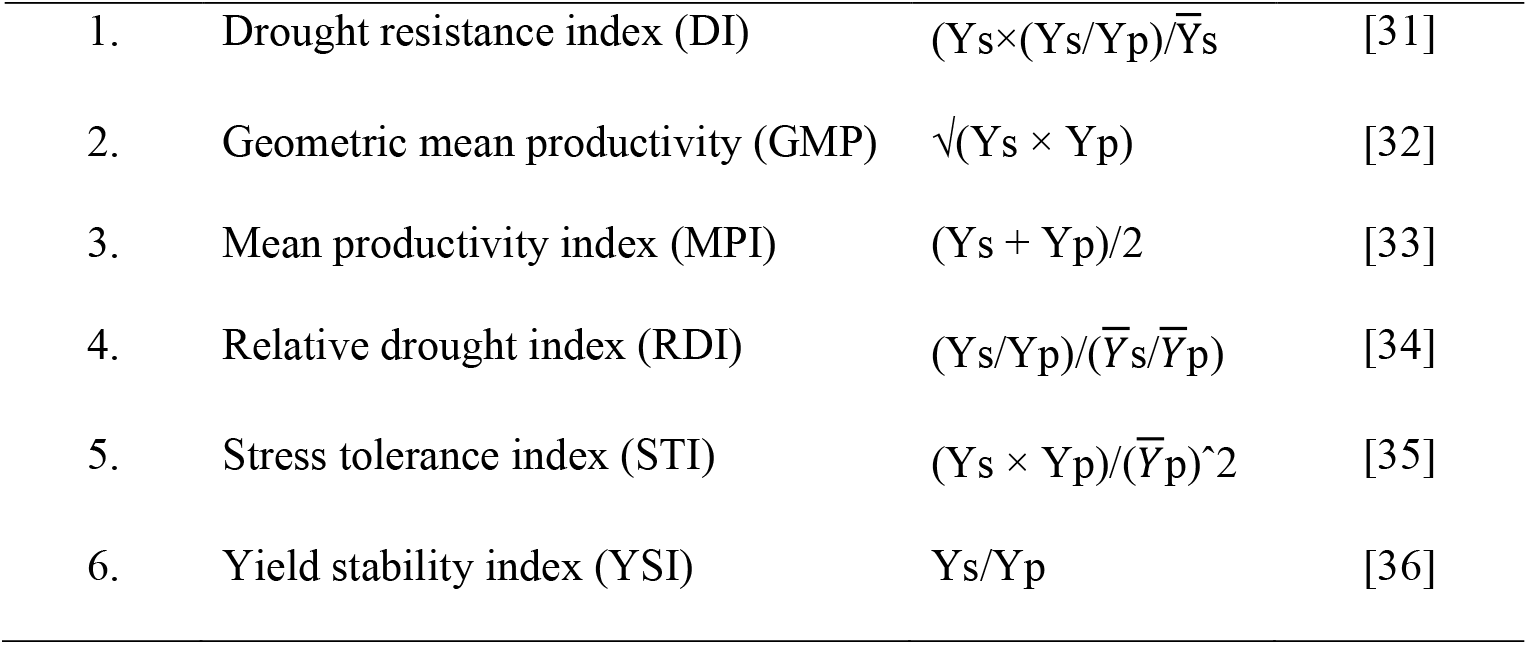

| Sl. No. | Stress indices | Formula | Reference |
| --- | --- | --- | --- |
| 1. | Drought resistance index (DI) | $(Y_s \times (Y_s/Y_p))/\bar{Y}_s$ | [31] |
| 2. | Geometric mean productivity (GMP) | $\sqrt{(Y_s \times Y_p)}$ | [32] |
| 3. | Mean productivity index (MPI) | $(Y_s + Y_p)/2$ | [33] |
| 4. | Relative drought index (RDI) | $(Y_s/Y_p)/(\bar{Y}_s/\bar{Y}_p)$ | [34] |
| 5. | Stress tolerance index (STI) | $(Y_s \times Y_p)/(\bar{Y}_p)^2$ | [35] |
| 6. | Yield stability index (YSI) | $Y_s/Y_p$ | [36] |

Where Ys and 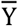s is the trait value and mean of the trait under drought treatment, respectively and Yp and 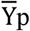 is the value for the same trait and mean value under the control condition, respectively.

### 2.4 Statistical analysis

#### 2.4.1 Analysis of variance

Analysis of variance was done after confirming the normal distribution of the data by performing skewness, kurtosis and Shapiro Wilk test at a 5% significance level.

#### 2.4.2 Multiplicative interactive AMMI ANOVA

Traits under the present study were subjected to ANOVA following the AMMI model [37,38]. Genotypes performing stable in each environment were chosen by AMMI analysis and tested for statistical significance by Gollob’s (1968) F-test procedure [39]. AMMI bi-plots were constructed using mean grain yield and principal component analysis (PCA). ANOVA and AMMI ANOVA were performed using Genstat 18^th^ edition (32-bit). Accuracy in the prediction of AMMI and BLUP models was tested by comparing RMSPD (Root Mean Square Prediction Difference) estimates of the models used [40].

Gauch and Zobel’s (1998) AMMI equation is as follows:

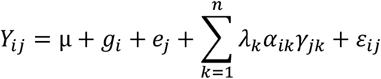

Where,

Y_ij_ is the grain yield value of i^th^ genotype in the j^th^ Environment, µ is the mean grain yield value, g_i_ and ej are the means deviations of i^th^ genotype, and j^th^ environment from the mean grain yield, respectively. Λ_k_ is the square root of the eigen value of the k^th^ IPC axis, α_ik_ and γ_jk_ are the interaction IPC scores for k^th^ IPC of the i^th^ genotype and j^th^ environment, respectively, and ε_ij_ is the residual.

To identify specific/broad adaptation of genotypes across environments, the AMMI stability value (ASV) was estimated [41].

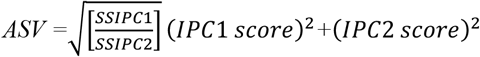

Where,

SSIPC 1 and SSIPC 2 are the sums of squares attributable to the first two interaction principal components (IPC1 and IPC2). ASV is the distance from the origin of the 2D scatter diagram of IPCA 1 *vs* IPCA 2 scores [41]. Lower ASV values represent the more excellent stability of a genotype across environments.

#### 2.4.3 GGE bi-plot analysis

GGE bi-plot analysis combines the effects of genotype and genotype × environment interaction [42,43]. GGE biplots were analyzed using GEA-R software (CIMMYT, 2015). The model of the GGE bi-plot is:

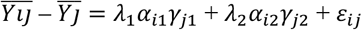

Where,

Y_ij_ = mean grain yield of i^th^ genotype in the j^th^ environment; 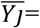 mean grain yield of all the genotypes in the j^th^ environment; λ_1_ and λ_2_ are the square roots of Eigen values of first and second genotype-by-environment interaction principal components (IPC) axes, respectively; α_i1_ and α_i2_ are the scores of the first and second IPC respectively for the i^th^ genotype, γ_i1_ and γ_i2_ are the first and second IPCs respectively for the i^th^ environment, and ε_ij_ is the residual.

#### 2.4.4 Multi-trait stability index (MTSI) analysis

The stability of each genotype across environments (GEI) was estimated by WAASB (Weighted Average of Absolute score obtained by Singular value decomposition from the Best linear unbiased predictor using linear mixed effect model). Further, a multi-trait stability index (MTSI) was also performed [44]. MTSI and WASSB analyses were performed in the metan [45] R package. The equation for MTSI is as follows:

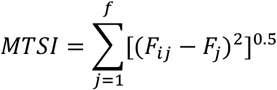

Where,

MTSI is the multi-trait stability index for the **i**^th^ genotype, **F**_**ij**_ is the **j**^th^ score of the **i**^th^ genotype, and **F**_**j**_ is the **j**^th^ score of the ideotype.

## 3. RESULTS

### 3.1 Analysis of variance

Data from all the environments were under the normal distribution. ANOVA was performed and the total variation was partitioned between genotype, environment, genotype × environment interactions and error. The mean sum of squares of all the traits significantly differed among genotype, environment and genotype × environment interactions (Table 1).

**Table 1.**
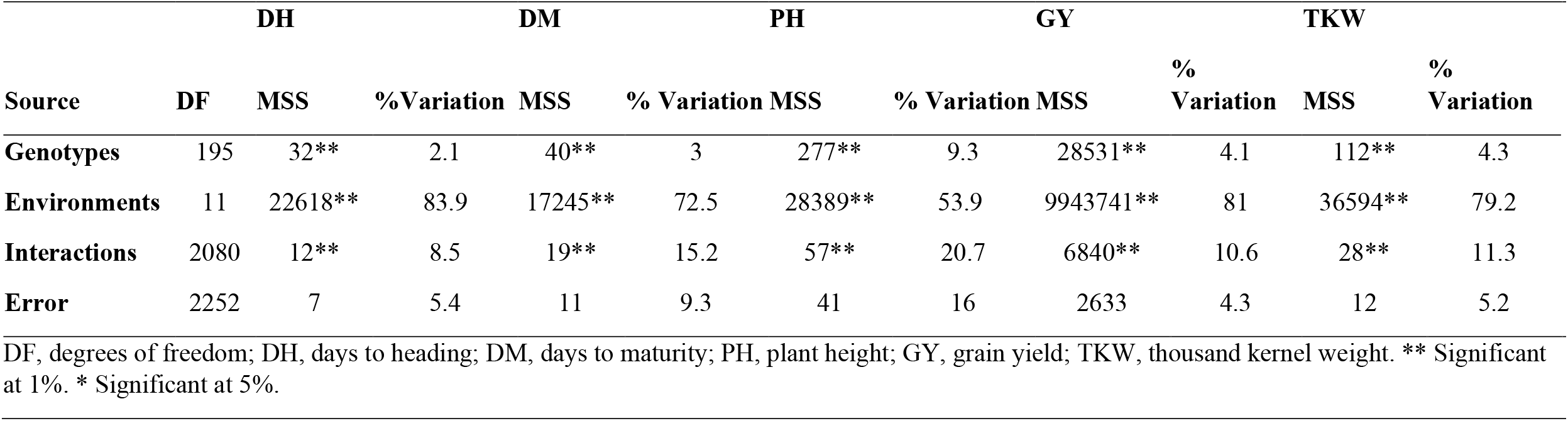
ANOVA for traits under investigation of 196 genotypes under control and drought treatment during *Rabi* 2020-21 and 2021-22.

The percent variation attributed towards total variability (G + E + G×E) is maximum for DH (83.9%) followed by GY (81%), TKW (79.2.6%), DM (72.5%) and PH (53.9%). Variation attributed by genotype was about 9.3% for PH followed by TKW (4.3%), GY (4.1%), DM (3%) and DH (2.1%). The interaction between genotype with environment (GEI) reported higher for PH (20.7%) followed by DM (15.2%), TKW (11.3%), GY (10.6%) and DH (8.5%) (Table 1).

AMMI ANOVA performed for grain yield partitioned total variability into genotype, environment and interaction (GEI). The environment contributed a higher variation (81%), followed by interaction (10.6%) and genotype (4.1%). The interaction component was further split into 5 interaction principal components where IPCA1 contributed 36.5% variation followed by IPCA2 (23.7%), IPCA3 (15.6%), IPCA4 (7.5%) and IPCA5 (4.9%) (Table 2).

**Table 2.** AMMI analysis of variance for grain yield among 196 genotypes of wheat under 12 environments during *Rabi* 2020-21 and 2021-22.

| Source | DF | SS | MSS | % Variation | GEI Contribution explained (%) |
| --- | --- | --- | --- | --- | --- |
| **Genotypes** | 195 | 5563460 | 28531** | 4.1 |  |
| **Environments** | 11 | 109381156 | 9943741** | 81 |  |
| **Interactions** | 2085 | 14260814 | 6840** | 10.6 |  |
| **IPCA 1** | 205 | 5207229 | 25401 |  | 36.5 |
| **IPCA 2** | 203 | 3382705 | 16664 |  | 23.7 |
| **IPCA 3** | 201 | 2224249 | 11066 |  | 15.6 |
| **IPCA 4** | 199 | 1069421 | 5374 |  | 7.5 |
| **IPCA 5** | 197 | 694804 | 3527 |  | 4.9 |
| **Error** | 2224 | 5856025 | 2633 | 4.3 |  |

DF, degrees of freedom; SS, sum of squares; MSS, mean sum of squares; GEI, genotype environment interaction; IPCA 1, interaction principal component 1; IPCA 2, interaction principal component 2; IPCA 3, interaction principal component 3; IPCA 4, interaction principal component 4; IPCA 5, interaction principal component 5. \*\* Significant at 1%.

### 3.2 Correlation study of grain yield and stress tolerance index

Under all the tested environments, grain yield under control and grain yield under drought treatment showed positive correlation (Fig. 2) (during 2020-21 trial at BHU, (r= 0.53**), at Pune (r= 0.44**) and at Jabalpur (r= 0.43**)., and during 2021-22 trial at BHU, (r= 0.57**), at Pune (r= 0.53**) and at Jabalpur (r= 0.43**). We further analyzed various stress indices (mentioned in materials and methods) to evaluate the performance of wheat genotypes under drought stress and calculated the correlation among these indices.

**Fig. 2.**
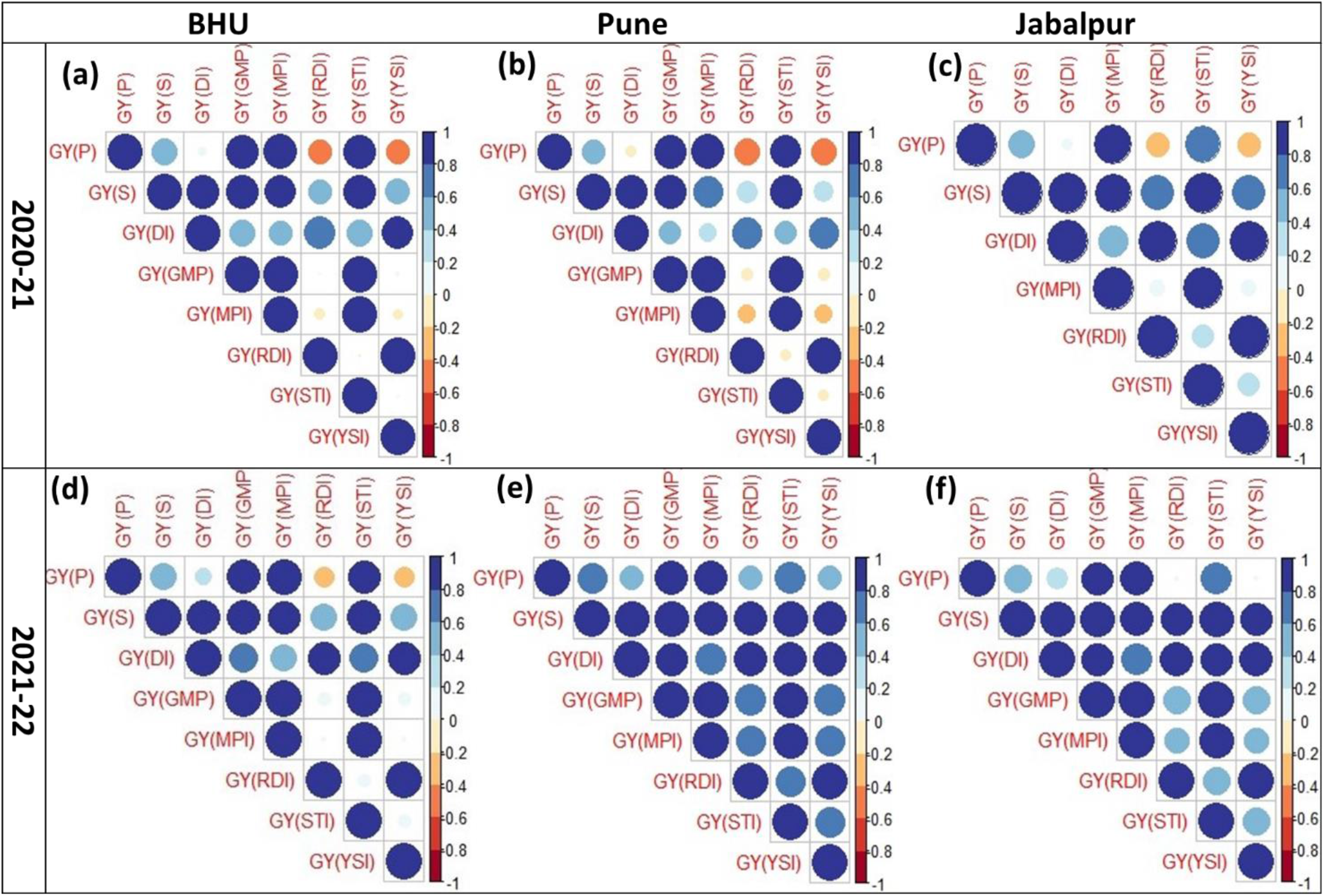
Correlation analysis between stress indices and grain yield under irrigated control and drought treatment at BHU during 2020-21 (a) and 2021-22 (d), at Pune during 2020-21(b) and 2021-22 (e), and at Jabalpur during 2020-21 (c) and 2021-22 (f). GY(P): grain yield under control; GY (S): grain yield under stress, GY(DI): Drought resistance index, GY(GMP): Geometric mean productivity, GY(MPI): Mean productivity index, GY(RDI): Relative drought index, GY(STI): Stress tolerance index, GY(YI): Yield index, GY(YSI): Yield stability index.

At BHU location, grain yield under drought had positive and significant correlation with stress indices: DI (r = 0.87**), GMP (r = 0.89**), MPI (r = 0.83**), STI (r = 0.88**) during 2020-2021 trial (Fig. 2a), and DI (r = 0.91**), GMP (r = 0.89**), MPI (r = 0.82**) and STI (r = 0.89**) during 2021-22 trial (Fig. 2d). Drought tolerant and susceptible genotypes were classified based on the stress indices with a higher correlation coefficient with the grain yield under drought during 2020-21 and 2021-22. Genotypes were ranked based on stress indices and with the mean rank of two consecutive years. The extreme 30 genotypes (top and bottom rank) are presented in Supplementary Table 2.

At Pune location, grain yield under drought treatment showed highly positive and significant correlation with stress indices DI (r = 0.84**), GMP (r = 0.85**), MPI (r = 0.73*), STI (r = 0.84**) during 2020-2021 trial (Fig. 2b) and DI (r = 0.96**), GMP (r = 0.94**), MPI (r = 0.90**), STI (r = 0.94**) during 2021-2022 trial (Fig. 2e). The extreme 30 genotypes are presented in the SupplementaryTable 2.

At Jabalpur location, grain yield under drought treatment reported highly positive and significant correlation with DI (r = 0.92**), MPI (r = 0.81**), STI (r = 0.89**) during 2020-2021 trial (Fig. 2c), and DI (r = 0.96**), MPI (r = 0.89**), STI (r = 0.94**) during 2021-2022 trial (Fig. 2f). The extreme 30 genotypes are presented in the Supplementary Table 2.

### 3.3 Correlation of grain yield with NDVI, SPAD and CTD

Physiological parameters such as NDVI, SPAD and CTD were recorded at the BHU center for two consecutive years 2020-21 and 2021-22. Whereas NDVI and CTD were recorded only during *Rabi* 2021-22 at Pune and Jabalpur, correlation analysis was performed among these traits at different developmental stages (heading, grain filling and maturity).

At the BHU center, NDVI showed no or poor correlation with grain yield under control (Fig. 3). However, NDVI recorded a positive and significant correlation (r = 0.41** 2020-21, r = 0.36** 2021-22) (Fig. 3) with grain yield under drought at the maturity stage. At Pune and Jabalpur centers also NDVI recorded a significantly higher correlation with grain yield at maturity under drought (Pune: r = 0.39** 2021-22; Jabalpur: r = 0.33** 2021-22). Control treatments had a less but significant correlation between NDVI and grain yield (Pune: r = 0.17* 2021-22; Jabalpur: r = 0.18* 2021-22) (Fig. 3). Other stages (heading and grain filling stages) also had a significantly positive correlation between NDVI and grain yield under drought at, both, Pune and Jabalpur locations (Fig. 3). During maturity, NDVI readings ranged between 69.5 to 78.0; and 31.0 to 63.5 with a mean of 73.3 and 47.6 under control in consecutive two years; however, under drought, it ranged from 19.5 to 57.5 and 14.0 to 45.0 with a mean of 35.3 and 26.8 in consecutive two years at BHU. The detailed observations of NDVI recorded at different stages and locations were provided in Supplementary Table 3a-d.

**Fig. 3.**
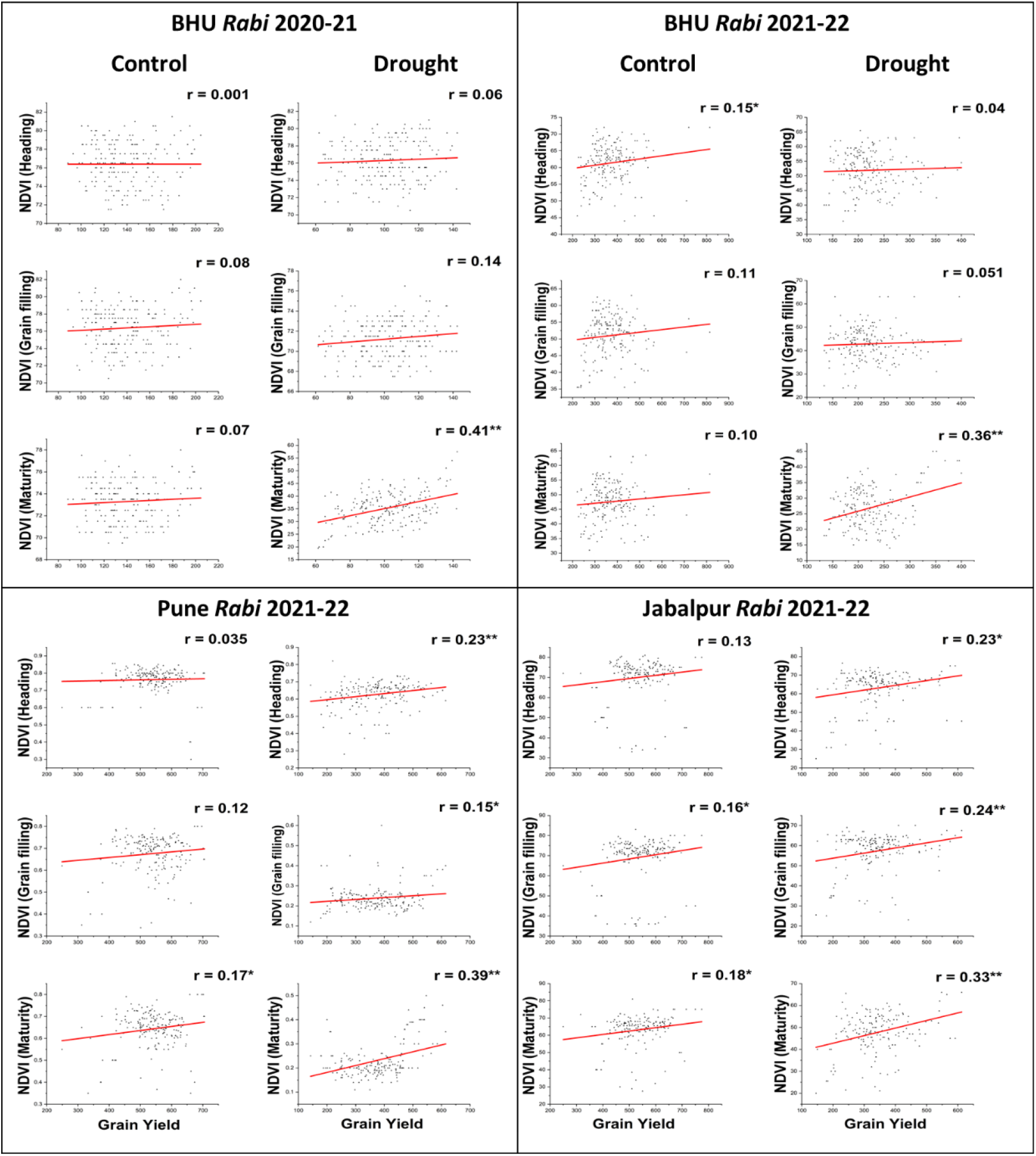
Regression analysis between grain yield vs NDVI under control and drought treatment during heading, grain filling and maturity at three locations BHU, Pune and Jabalpur. * and ** represent significance at 5% and 1% probability levels, respectively. Values without * represent not significant.

SPAD also showed a statistically significant correlation with grain yield at maturity for both, control (r = 0.36**) and drought (r = 0.45**) treatments (Supplementary Fig. 1). No significant correlations were observed for the other two growth stages (Supplementary Fig. 1). SPAD values recorded were analyzed for mean, range and standard deviation, same were detailed in Supplementary Table 3e.

At all developmental stages, CTD reported a poor correlation with grain yield under control and drought treatment at all locations (Supplementary Fig. 2). Detailed readings of mean, standard deviation and range recorded at all locations been presented in Supplementary Table 3a-d.

### 3.4 AMMI Bi-plot

AMMI Bi-plot was plotted considering grain yield and IPC1 (Interaction Principal Component 1) scores. The degree of environmental influence on genotypes was varied where E4 had the highest variation followed by E3, E6, E7, E11, E9, E10, E5, E12, E8, E1 and E2. The interaction component IPCA1 attributed 43.43% of the total variation with grain yield (Fig. 4a).

**Fig. 4.**
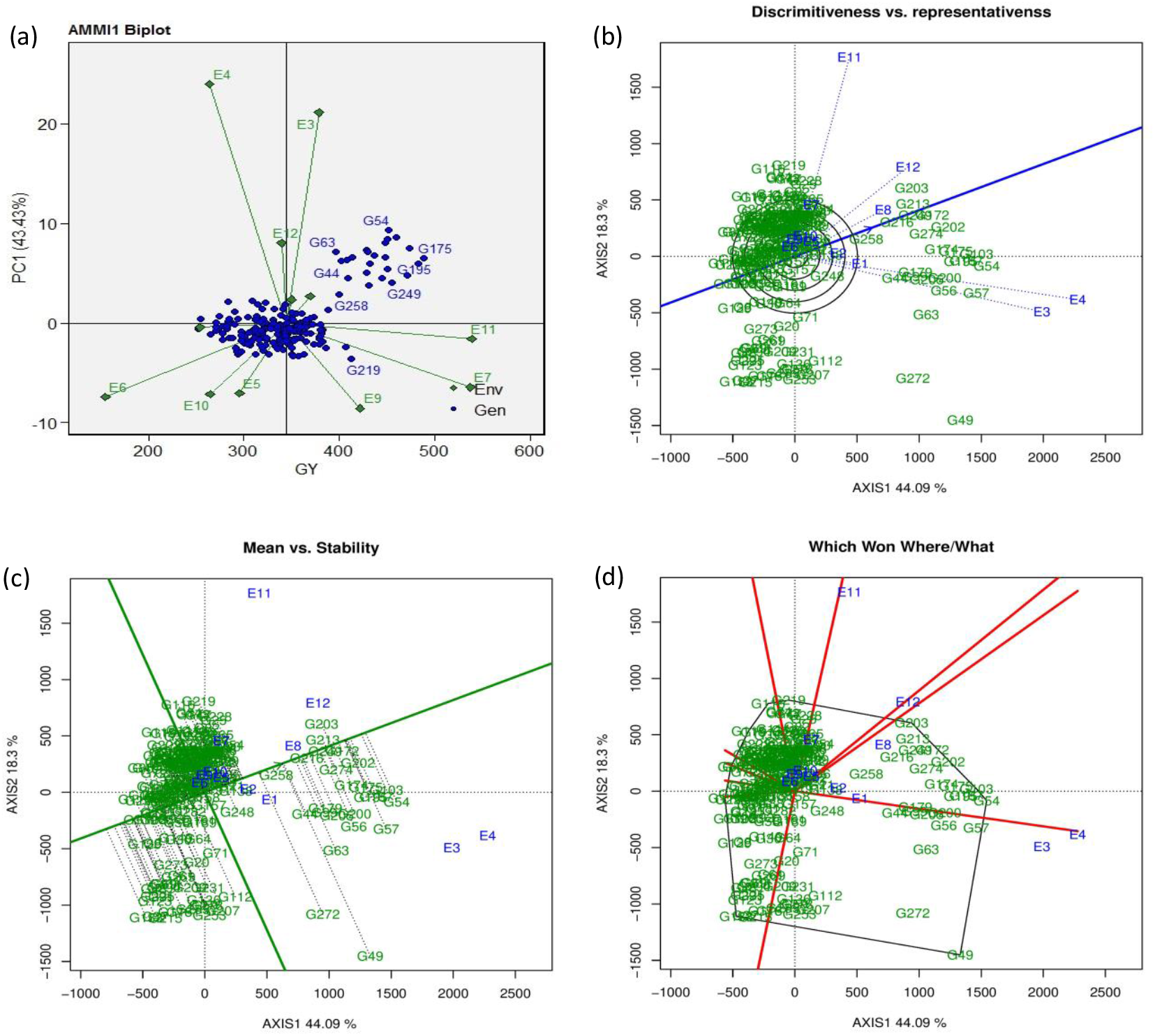
(a) AMMI1 Bi-plot (grain yield vs IPC1), (b) discrimination and representativeness view of GGE bi-plots, (c) Mean vs Stability view of GGE bi-plot, (d) Polygon view of the GGE bi-plot for grain yield of 196 genotypes and 12 environments under irrigated control and drought treatment during *Rabi* 2020-21 and 2021-22.

The stability index (SI) was analysed for genotypes under control and drought in all environments. Genotypes having lower SI were considered to be more stable. In the current study, G114, G280, G275, G289, G254, G269, G159, G177, G279, G112, G239, G236, G36, G264, G170, G252, G215, G82, G173, G291, G61, G46, G267, G262, G27, G206, G83, G231, G2 and G218 were extreme 30 genotypes that recorded the lowest SI in ascending order and, hence, considered more stable in all three locations under control and drought treatments. Whereas G150, G73, G148, G86, G248, G181, G119, G178, G257, G72, G110, G48, G283, G34, G87, G166, G118, G4, G169, G217, G161, G12, G151, G276, G149, G282, G221, G31, G8 and G32 were extreme 30 genotypes in descending order of SI values and considered least stable (Supplementary Table 4).

### 3.5 GGE bi-plots for grain yield trait

Graphical representations of GGE bi-plots were plotted considering principal interaction components (IPC1 and IPC2). The total variation attributed to IPC1 and IPC2 was 62.39% (Fig. 4b).

#### 3.5.1 Discriminative and representative view of GGE bi-plot

Among the tested environments, E4, E3 and E11 represented the most discriminative, whereas E6, E9 and E10 were the least discriminative environments. The angel between the E3, E4, E1, E2, E5, E8, E12 and E10 were acute, indicating a strong positive correlation and similarities between these tested environments. E3 showed an obtuse angle with E6 and E9, and E4 showed an obtuse angle with E6 and E9, which suggested a negative correlation and cross-over interaction among these environments [46]. E3 and E4 observed right angles with E11 and E7 environments representing no relation. E8 represented the ideal test environment whose vector is closer to the average environment coordinate (AEC) for the grain yield (Fig. 4b).

#### 3.5.2 Mean vs stability view of GGE bi-plot

Genotypes G258, G216, G249, G274, G213, G172, G203, G174, G175, G254, G200, G275, G21, G138, G248, G23 and G236 were closer to the AEC arrow, with lower projections representing more stable performance in all the tested environments. However, genotypes G133, G37, G123, G215, G225, G48, G178 and G151, located opposite the AEC coordinate for grain yield trait, indicated the least stability (Fig. 4c).

#### 3.5.3 Which one where the view of the GGE bi-plot

We further determined the best performing genotypes for target-specific locations. The GGE bi-plot was classified into 10 mega environments. E3 and E4; E1, E2 and E8; E12; E11, E7, E10 and E5; E9, and E6 fall separately under six different mega environments. G21 performed superior in the E1 environment, G57 was the best performer at E2 and E4, G54 at E3, G32 at E5, G269 at E6, G47 at E7 and E8, G272 at E9 and E10, G116 at E11and G175 at E12 (Fig. 4d).

### 3.6 Selection of genotypes based on multi-trait stability index (MTSI)

The likelihood ratio test (LRT) reported highly significant (p < 0.05) for genotype and GEI effect (Supplementary Table 5). Pearson’s correlation matrix for WAASBY (a combination of WAASB Score and mean grain yield) values was estimated, and highly associated traits were grouped under a common factor. An exploratory factor analysis retained three principal components (FA1, FA2 and FA3), contributing 75.22% of the total variance (Supplementary Table 6). Communality depicts the proportion of each trait variance, explained by factor analysis. Communality reported a maximum for the GY (0.93) and a minimum for the PH (0.62) with an average communality (h) of 0.75. Identifying stable and high mean performers under MTSI is based on the genotype-ideotype distance (Euclidian) using scores obtained under exploratory factor analysis. The genotype scores and ideotype estimates for 196 genotypes obtained using explanatory factor analysis were given in Supplementary Table 7.

Selection differential (SD%) of WAASB scores were positive for all traits, inferring the method was effective in selecting high mean performing and most stable genotypes. PH (15.5 %) and GY (14.4 %) reported highest selection differential under WAASBY index and DH reported the lowest (9.2 %) (Table 3).

**Table 3.**
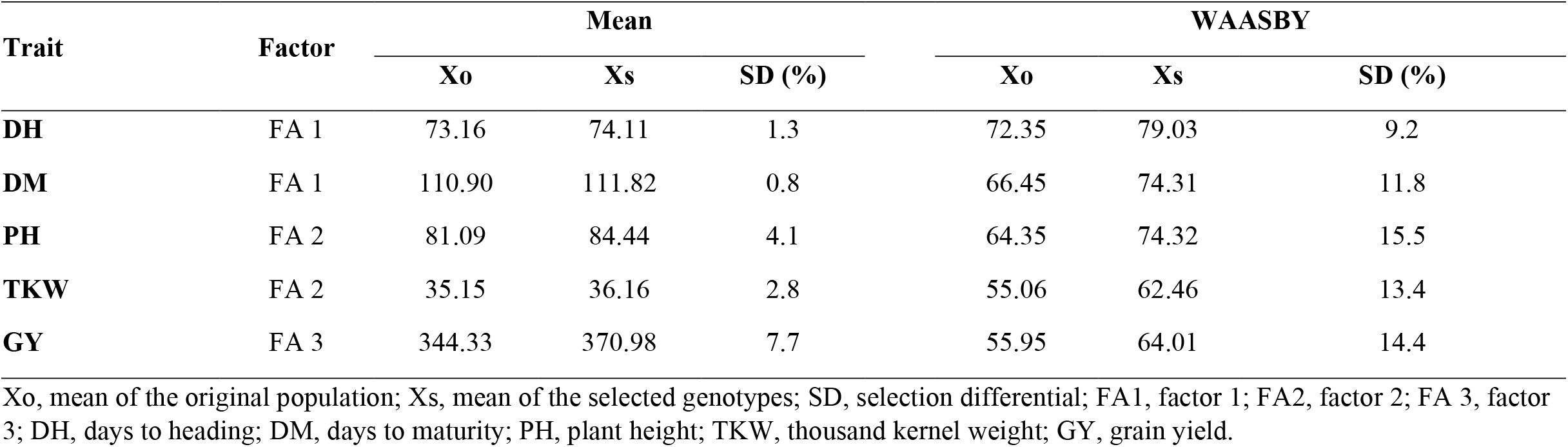
Selection differential (SD) in percent for mean performance and WAASBY index for 5 traits of 196 genotypes across twelve environments during *Rabi* 2020-21 and 2021-22.

Y × WAASB bi-plot represented mean grain yield at the x-axis and WAASB values at the y-axis. Bi-plot analysis plotted E1, E2, E4, E5, E6, E10 and E12 environments under the first quadrant and E3, E7, E8, E9 and E11 under the second quadrant. The genotypes were classified into four quadrants where genotypes falling under the first quadrant were unstable and low yielding, the second quadrant was unstable but had a mean above the grand mean, and the third quadrant was stable but low yield. In contrast, the fourth quadrant represents more stable and highly productive genotypes (Fig. 5). Sixty-three genotypes fell under the fourth quadrant, representing more stable and high mean performance than other genotypes (Supplementary Table 8).

**Fig. 5.**
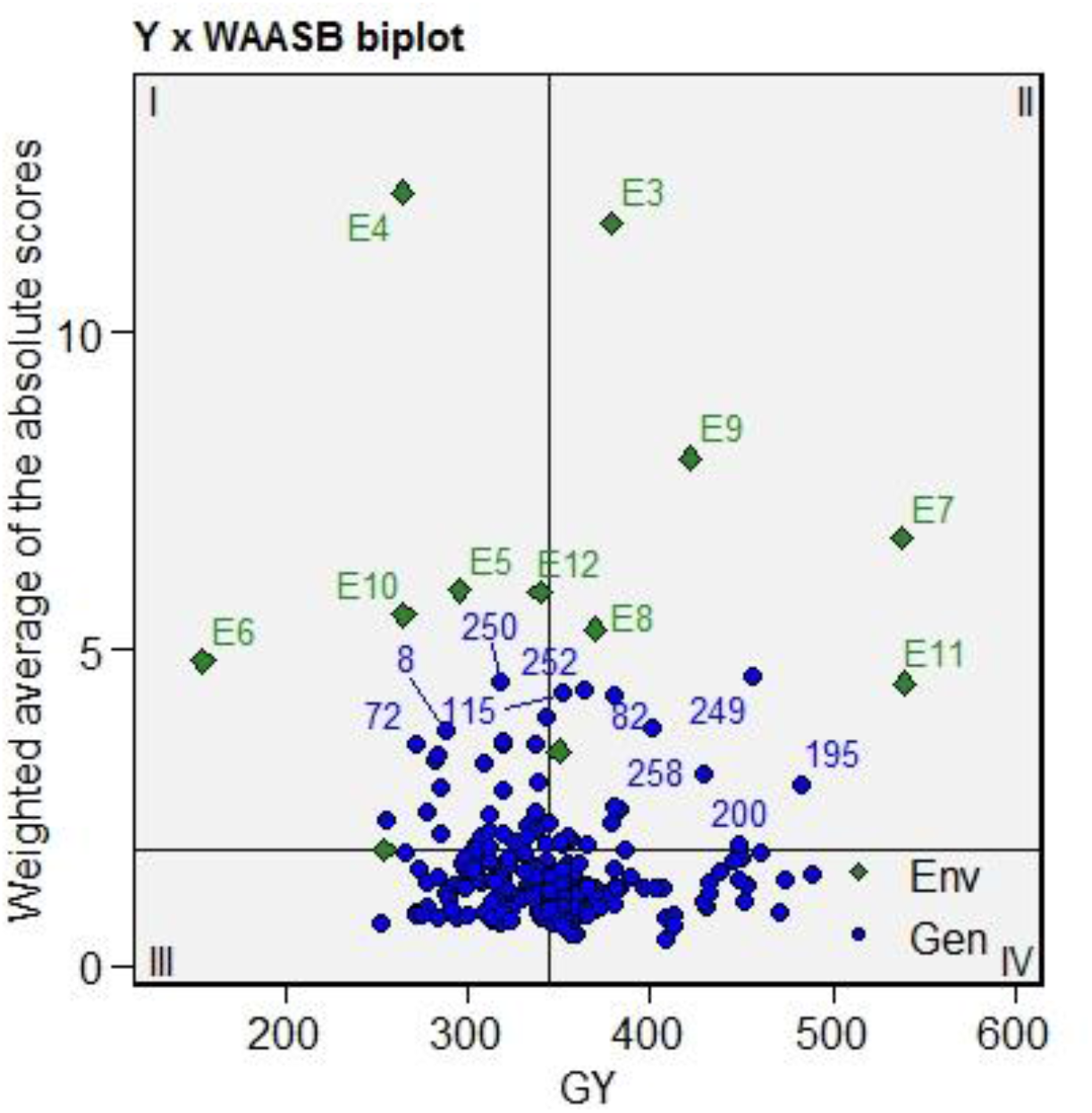
Yield × WAASB bi-plot based on joint interpretation of productivity (Y) and stability (WAASB) for 196 genotypes evaluated under 12 environments during *Rabi* 2020-21 and 2021-22.

Lowering the MTSI index indicated higher genotypic stability and high mean performance [44]. MTSI was estimated by considering DH, DM, PH, TKW and GY traits in the present study. Genotypes were selected with the selection intensity of 15%, as represented in Fig 6 and Supplementary Table 7. 29 Genotypes (G249, G281, G173, G169, G192, G213, G175, G21, G209, G275, G86, G84, G267, G228, G276, G103, G158, G286, G45, G241, G188, G187, G198, G212, G191, G126, G232, G172 and G113) located out of the red circle indicated higher adaptability to all environments and were considered as best performing genotypes. Selection intensity is considered a cut-off MTSI value of 5.42. Genotypes such as G108, G74, G231, G88, G8, G293, G294, G136, G30, G129, G23, G105, G37, G69, G89, G35, G27, G112, G26 and G96 were positioned towards the origin with black circles, indicating poor performance and stability.

**Fig. 6.**
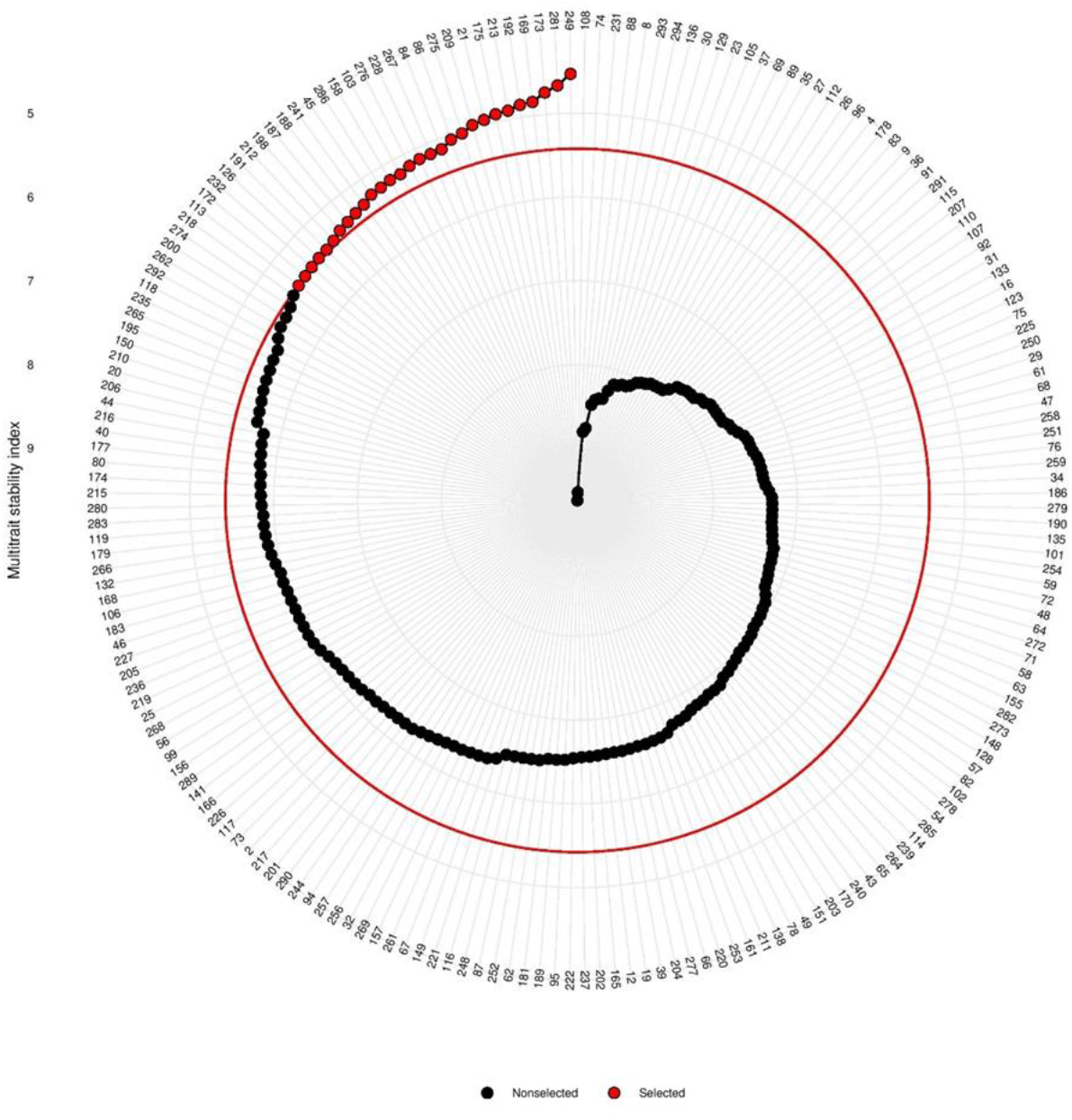
Genotype ranking and selected genotypes for the multi-trait stability index considering a selection intensity of 15 % for 196 genotypes evaluated under 12 environments during *Rabi* 2020-21 and 2021-22.

### 3.7 Testing the accuracy of models

RMSPD test was performed to predict the accuracy of AMMI and BLUP-based models for grain yield trait. A model with a lower RMSPD value is regarded as the most accurate and vice-versa. The means of each model are based on 50 estimates of RMSPD (Fig. 7 and Supplementary Table 9). RMSPD of our study depicted the BLUP-based model (RMSPD = 70.5) as the most accurate as compared to AMMI models. WAASBY and MTSI models use BLUP to classify the most stable and least stable genotypes. The stable genotypes selected using WAASBY and MTSI models showed a mean greater than the overall mean of all genotypes.

**Fig. 7.**
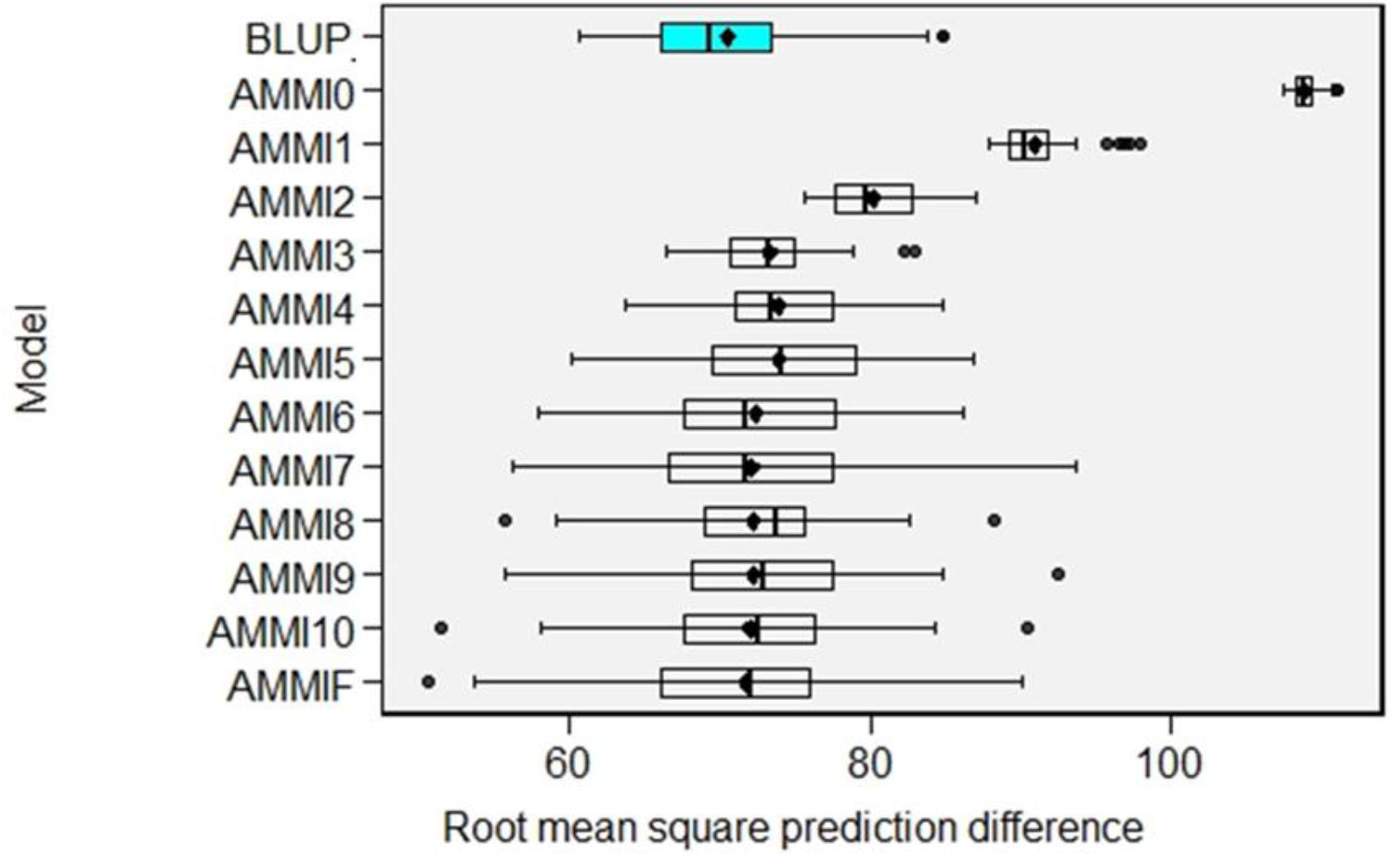
Box plots indicating RMSPD values of BLUP and AMMI models tested for grain yield.

## 5. Discussion

The development of drought-tolerant cultivars performing stably across environments are of prime importance in the changing climatic scenario of present-day agriculture. To meet this objective, selecting drought-tolerant and stable cultivars is critical, which relies on appropriate statistical models. Therefore, understanding drought adaptation mechanisms and evaluating diverse breeding material in multi-environmental locations with novel statistical models to select the stable lines under drought is very much necessary [47]. The present study observed significant genetic variability for drought tolerance in all genotypes, indicating the diverse nature of the wheat population. In addition, all three geographical locations (Varanasi, Pune and Jabalpur) showed high variability as revealed by poor correlation among different sites under control as well drought treatments, suggesting a significant contribution from GEI to all drought-associated traits. Days to heading and grain yield were identified as highly influenced traits in different environments. Grain yield had the highest contribution from GEI than the genotype inferring the complexity of selecting genotypes performing stable and widely adaptable across varied environments [48]. Overall, genotypic variation and environmental conditions cited in the present study served to understand drought adaptation mechanisms in wheat.

Identifying ideal environments to select desirable genotypes for farmers is the prime objective of every breeding programme [24,49]. The present study identified E8 as an ideal environment for selecting genotypes with enhanced drought tolerance. Further, Bi-plot analysis revealed a few genotypes performing stable across different environments or in a specific environment. Mean v/s stability bi-plot analysis has previously been used to identify best performing genotypes across environments [50–53]; however, Which Won Where bi-plot provides superior genotypes for a specific environment [54,55].

Drought stress adversely affects plant growth by accelerating senescence, lowering photosynthesis and producing poor yield [18]. The maintenance of greenness allows a genotype to prolong the photosynthesis and the grain filling duration, which may provide a yield advantage especially under severe water deficit conditions [18,56]. We observed a positive and significant correlation of NDVI with grain yield specifically under drought but not under control at the BHU center. In addition, this correlation was specific to the maturity stage and not to other stages (heading and grain filling). A positive correlation between NDVI with grain yield was reported for heat and combined effects of heat and drought; however, drought had no significant correlation between NDVI and grain yield at the stage of physiological maturity [58]. An exact reason is not clear but differences in imposing water stress treatment and environmental conditions might be the possible reasons for data discrepancy between the two studies. Overall, a specific correlation between NDVI and grain yield under drought may provide opportunities for breeders to use the NDVI as a potential screening tool to select drought tolerant genotypes under water stress conditions in wheat.

Previous studies used grain yield as primary criteria, and the contribution of other important traits in adaptation to different environmental conditions was not considered in selecting stable genotypes [59,60]. However, recently proposed linear mixed models WAASB and MTSI use both AMMI and BLUPs of all traits in screening genotypes across varied environments [44]. By utilizing WAASB and MTSI as a selection technique, genotypes performing well under water-limited conditions have been identified in maize [24], chickpea [26] and cassava [61]. We also found that WAASB and MTSI have reliable methods that identified stable and better-performing genotypes across environments. Further, MTSI results infer that the relative contribution (RC%) of the distance from genotype e.g., G218 (placed just below the cut point of selected stable genotypes in Fig. 6) to ideotype was ∼ 75% which is related to FA1 and FA2 (DH, DM, PH and TKW traits) (Supplementary Table 7). In contrast, the genotype G175 (stable genotype) has FA3 (GY) lowest relative contribution (1.4%) to MTSI, inferring G175 is close to the ideotype regarding grain yield trait. We propose further improving DH, DM, PH and TKW traits to enhance the stability of the lower-performing genotype (G218) across environments.

The stress indices used to evaluate the drought tolerance of wheat genotypes were found effective in screening wheat genotypes under drought. Our study identified reliable sources of drought tolerance with high mean performance for a specific environment as well as across multiple environments. The tolerant genotypes identified in this study are being utilized in the ongoing breeding program as donor parents to enhance the drought tolerance of wheat. In addition, these genotypes are important genetic resources to improve our understanding of drought adaptation mechanisms in wheat.

## 6. Conclusion

To identify novel sources of drought tolerance, we screened diverse wheat genotypes at different geographical locations under irrigated control and drought treatments. The data suggested NDVI at maturity as a reliable phenotypic marker in screening for drought tolerant genotypes under field conditions. Using various stress indices and mixed linear models such as MTSI and YxWASSB, a few genotypes were identified as drought tolerant and best performers across all tested environments. These genotypes are the potential genetic resources to understand drought adaptation mechanisms and be used as donor parents in breeding programmes to improve drought tolerance in wheat.

## Supporting information

"C:\Users\Bhu\Desktop\Lab students\Srinatha\Drought_WAMI\Two years data\supplementary tables final 17.9.22.zip"

## Acknowledgment

SS thanks DST-SERB project (M14-73) and seed money grant from IOE, BHU for financial support. SR acknowledge financial support obtained as a junior research fellow in the DST-SERB project (M14-73).

## Author contribution

SR, UK, PB, MV, SN and YKJ performed experiments; SR, GM, and RC analyzed the data and interpret results; SS and VKM planned, conceptualize, designed experiments and supervised the study; SR and SS wrote the initial draft and finalized it with VKM, RC, and AKJ. All authors approved the manuscript.

## Declaration

The authors declare no conflict of interest

## Abbreviations

WAMI: wheat associated mapping initiative
DH: days to heading
DM: days to maturity
PH: plant height
TKW: thousand kernel weight
GY: grain yield
NDVI: normalized difference vegetative index
SPAD: soil plant area development
CTD: canopy temperature depression
DI: drought resistance index
GMP: geometric mean productivity
MPI: mean productivity index
RDI: relative drought index
STI: stress tolerance index
YSI: yield stability index
Ys: yield under drought treatment;
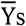: mean of the trait under drought treatment
Yp: yield under control condition;
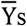: mean of the trait under control condition
GEI: genotype × environment interaction
AMMI: additive main effects, and multiplicative interaction
GGE: genotype main effect plus genotype-environment interaction effect
AEC: average-environment coordinate
SVD: singular value decomposition
ASV: AMMI stability value
YSI: yield stability index
IPCA: interaction principal component axis
RMSPD: root mean square prediction difference
BLUP: best linear unbiased prediction
LMM: linear mixed model
WAASB: a weighted average of absolute scores from the singular value decomposition of the matrix of BLUPs
MTSI: multi-trait stability index.

