## Supplementary material for "Spatio-temporal evaluation of drought adaptation in wheat revealed NDVI and MTSI as powerful tools for selecting tolerant genotypes": "C:\Users\Bhu\Desktop\Lab students\Srinatha\Drought_WAMI\Two years data\supplementary tables final 17.9.22.zip": supplementary tables final 17.9.22.docx

| Supplemental Table 1. Details of the test environments, treatment, season and location where 196 genotypes were evaluated during *Rabi* 2020-21 and 2021-22. | | | |
| --- | --- | --- | --- |
| Environment code | Treatment | Season | Location |
| E1 | Control | *Rabi* 2020-21 | BHU, Varanasi |
| E2 | Drought | *Rabi* 2020-21 | BHU, Varanasi |
| E3 | Control | *Rabi* 2021-22 | BHU, Varanasi |
| E4 | Drought | *Rabi* 2021-22 | BHU, Varanasi |
| E5 | Control | *Rabi* 2020-21 | ARI, Pune |
| E6 | Drought | *Rabi* 2020-21 | ARI, Pune |
| E7 | Control | *Rabi* 2021-22 | ARI, Pune |
| E8 | Drought | *Rabi* 2021-22 | ARI, Pune |
| E9 | Control | *Rabi* 2020-21 | BISA, Jabalpur |
| E10 | Drought | *Rabi* 2020-21 | BISA, Jabalpur |
| E11 | Control | *Rabi* 2021-22 | BISA, Jabalpur |
| E12 | Drought | *Rabi* 2021-22 | BISA, Jabalpur |

| Supplemental Table 2. List of selected 30 extreme genotypes based on stress indices and mean ranks in the two consecutive years 2020-21 and 2021-22 evaluated at 3 locations. | | | |
| --- | --- | --- | --- |
|  | BHU | Pune | Jabalpur |
| List top rank | G57, G195, G54, G187, G203, G49, G205, G200, G174, G272, G175, G216, G56, G172, G179, G99, G274, G138, G44, G103, G112, G202, G213, G249, G258, G231, G63, G71, G59 and G64 | G272, G39, G216, G200, G218, G49, G203, G19, G107, G269, G241, G273, G165, G115, G249, G195, G220, G228, G108, G25, G219, G47, G103, G172, G192, G40, G232, G266, G16 and G206 | G228, G80, G252, G206, G82, G218, G107, G251, G114, G103, G289, G39, G216, G195, G175, G179, G141, G249, G202, G126, G203, G138, G275, G219, G258, G254, G274 and G292 |
| Bottom rank | G40, G110, G221, G39, G16, G257, G148, G32, G96, G108, G35, G29, G128, G119, G86, G276, G129, G102, G106, G94, G72, G235, G4, G9, G250, G107, G26, G210, G191 and G34 | G133, G48, G265, G34, G237, G150, G129, G235, G29, G225, G253, G261, G166, G72, G91, G276, G169, G123, G257, G161, G248, G151, G148, G87, G73, G149, G114, G189 and G217 | G151, G35, G4, G231, G181, G9, G30, G72, G112, G253, G91, G61, G19, G209, G129, G123, G294, G235, G20, G285, G8, G133, G12, G59, G58, G26, G32, G119, G71 and G37 |

| Supplemental Table 3a. Mean, Standard deviation and range of NDVI, CTD reading and grain yield recorded under control and drought treatment at BHU *Rabi* 2020-21. GY( C): grain yield under control, GY(D): grain yield under drought, NDVIH: NDVI readings during heading, NDVIGF: NDVI readings during grain filling, NDVIM: NDVI readings during maturity, CTDH: CTD readings during heading, CTDGF: CTD readings during grain filling, CTDM: CTD readings during maturity, (C): under control, (D): under drought. | | | | | | | | | | | | | | |
| --- | --- | --- | --- | --- | --- | --- | --- | --- | --- | --- | --- | --- | --- | --- |
|  | GY  (C) | GY (D) | NDVIH (C) | NDVIH (D) | NDVIGF (C) | NDVIGF (D) | NDVIM (C) | NDVIM (D) | CTDH (C) | CTDH (D) | CTDGF (C) | CTDGF (D) | CTDM (C) | CTDM (D) |
| Mean | 139.8 | 101.5 | 76.4 | 76.3 | 76.4 | 71.2 | 73.3 | 35.3 | 11.7 | 7.0 | 11.6 | 6.9 | 8.7 | 3.9 |
| SD | 25.6 | 18.5 | 2.3 | 2.3 | 2.3 | 1.9 | 1.8 | 6.2 | 1.5 | 1.5 | 1.4 | 1.4 | 0.8 | 1.0 |
| Min | 88.5 | 61.5 | 71.5 | 70.5 | 70.5 | 67.5 | 69.5 | 19.5 | 8.5 | 3.5 | 8.4 | 3.0 | 6.3 | 1.5 |
| Max | 204.5 | 142.5 | 81.5 | 81.5 | 82.0 | 76.5 | 78.0 | 57.5 | 14.8 | 10.4 | 14.3 | 10.6 | 10.3 | 6.1 |

| Supplemental Table 3b. Mean, Standard deviation and range of NDVI, CTD reading and grain yield recorded under control and drought treatment at BHU *Rabi* 2021-22. | | | | | | | | | | | | | | |
| --- | --- | --- | --- | --- | --- | --- | --- | --- | --- | --- | --- | --- | --- | --- |
|  | GY (C) | GY (D) | NDVIH (C) | NDVIH (D) | NDVIGF (C) | NDVIGF (D) | NDVIM (C) | NDVIM (D) | CTH (C) | CTH (D) | CTGF (C) | CTGF (D) | CTM (C) | CTM (D) |
| MEAN | 371.3 | 222.4 | 61.3 | 51.9 | 51.0 | 42.9 | 47.6 | 26.8 | 8.4 | 6.3 | 10.2 | 6.4 | 12.3 | 6.3 |
| SD | 86.6 | 55.0 | 5.6 | 6.3 | 6.0 | 6.9 | 6.1 | 6.3 | 1.9 | 2.0 | 1.8 | 2.0 | 2.1 | 1.7 |
| MIN | 221.0 | 133.7 | 44.0 | 34.5 | 35.5 | 24.0 | 31.0 | 14.0 | 3.4 | 1.4 | 5.1 | 2.1 | 6.4 | 2.0 |
| MAX | 814.4 | 514.7 | 72.0 | 65.5 | 63.0 | 63.0 | 63.5 | 45.0 | 12.1 | 11.3 | 13.8 | 11.3 | 16.8 | 9.6 |

| Supplemental Table 3c. Mean, Standard deviation and range of grain yield and physiological parameters recorded under irrigated control and drought treatment during heading, grain filling and maturity at Pune *Rabi* 2021-22. | | | | | | | | | | | | | | |
| --- | --- | --- | --- | --- | --- | --- | --- | --- | --- | --- | --- | --- | --- | --- |
|  | GY (C) | GY (D) | NDVIH (C) | NDVIH (D) | NDVIGF (C) | NDVIGF (D) | NDVIM (C) | NDVIM (D) | CTDH (C) | CTDH (D) | CTDGF (C) | CTDGF (D) | CTDM (C) | CTDM (D) |
| Mean | 541.97 | 370.36 | 0.77 | 0.68 | 0.71 | 0.60 | 0.20 | 0.20 | 8.86 | 5.21 | 10.12 | 6.91 | 9.72 | 8.42 |
| SD | 61.74 | 76.17 | 0.04 | 0.06 | 0.04 | 0.08 | 0.04 | 0.03 | 1.86 | 2.15 | 1.64 | 1.20 | 1.37 | 2.18 |
| Min | 392.00 | 213.00 | 0.63 | 0.47 | 0.58 | 0.07 | 0.10 | 0.15 | 1.10 | -0.10 | 4.55 | 4.60 | 4.72 | 3.90 |
| Max | 683.00 | 517.50 | 0.86 | 0.82 | 0.79 | 0.75 | 0.30 | 0.28 | 13.65 | 10.10 | 14.60 | 9.90 | 13.70 | 12.42 |

| Supplemental Table 3d. Mean, Standard deviation and range of grain yield and physiological parameters recorded under irrigated control and drought treatment during heading, grain filling and maturity at Jabalpur *Rabi* 2021-22. | | | | | | | | | | | | | | |
| --- | --- | --- | --- | --- | --- | --- | --- | --- | --- | --- | --- | --- | --- | --- |
|  | GY (C) | GY (S) | NDVIH (C) | NDVIH (D) | NDVIA (C) | NDVIA (D) | NDVIG (C) | NDVIM (D) | CTDH (C) | CTDH (D) | CTDGF (C) | CTDGF (D) | CTM (C) | CTM (D) |
| Mean | 538.9 | 339.7 | 70.1 | 62.8 | 69.2 | 57.4 | 63.2 | 47.7 | 6.5 | 7.0 | 7.8 | 7.0 | 5.2 | 6.4 |
| SD | 82.8 | 89.1 | 9.9 | 10.0 | 11.0 | 9.5 | 9.0 | 9.3 | 1.1 | 1.0 | 1.2 | 0.9 | 1.2 | 1.1 |
| Min | 250.0 | 146.0 | 33.0 | 25.0 | 35.0 | 23.0 | 27.5 | 20.0 | 3.4 | 5.0 | 4.6 | 3.7 | 3.0 | 3.8 |
| Max | 774.0 | 612.5 | 81.0 | 76.5 | 83.0 | 70.0 | 81.0 | 66.0 | 9.4 | 9.7 | 11.0 | 10.0 | 8.2 | 10.4 |

| Supplemental Table 3e. Mean, Standard deviation and range of SPAD reading and grain yield recorded under control and drought treatment at BHU during *Rabi* 2020-21 and 2021-22. GY( C): grain yield under control, GY(D): grain yield under drought, SPADH: SPAD readings during heading, SPADGF: SPAD readings during grain filling, SPADM: SPAD readings during maturity, (C): under control, (D): under drought. | | | | | | | | | | | | | | | | | |
| --- | --- | --- | --- | --- | --- | --- | --- | --- | --- | --- | --- | --- | --- | --- | --- | --- | --- |
|  | at BHU *Rabi* 2020-21 | | | | | | | | | at BHU *Rabi* 2021-22 | | | | | | | |
|  | GY (C) | GY (D) | SPADH (C) | SPADH (D) | SPADGF (C) | SPADGF (D) | SPADM (C) | SPADM (D) | GY (C) | | GY (D) | SPADH (C) | SPADH (D) | SPADGF (C) | SPADGF (D) | SPADM (C) | SPADM (D) |
| Mean | 139.8 | 101.5 | 47.8 | 42.7 | 49.5 | 49.9 | 47.4 | 42.4 | 371.3 | | 222.4 | 45.2 | 45.1 | 47.2 | 42.8 | 44.9 | 44.2 |
| SD | 25.6 | 18.5 | 2.7 | 3.3 | 2.8 | 2.6 | 3.0 | 2.9 | 86.6 | | 55.0 | 3.6 | 3.9 | 3.9 | 2.9 | 4.0 | 3.4 |
| Min | 88.5 | 61.5 | 41.5 | 34.5 | 43.4 | 44.0 | 40.7 | 34.9 | 221.0 | | 133.7 | 35.5 | 34.5 | 38.0 | 37.0 | 37.8 | 35.0 |
| Max | 204.5 | 142.5 | 56.0 | 49.3 | 57.8 | 58.3 | 56.2 | 50.3 | 814.4 | | 514.7 | 60.0 | 57.0 | 60.0 | 52.0 | 58.0 | 54.0 |

| Supplemental Table 4. Grain yield, AMMI stability values (ASV), multi-trait stability index (MTSI) and ranking of extreme 30 highly stable and less stable genotypes tested across twelve environments during *Rabi* 2020–21 and 2021-22. | | | | | | | | | |
| --- | --- | --- | --- | --- | --- | --- | --- | --- | --- |
| **Genotype** | **Mean** | **rMean (A)** | **IPCAg1** | **IPCAg2** | **ASV** | **rASI(B)** | **SI (A+B)** | **MTSI** | **rMTSI** |
| **Highly Stable** | |  |  |  |  |  |  |  |  |
| G114 | 275.3 | 40 | 0.11 | -0.14 | 0.19 | 2 | 42 | 7.6 | 118 |
| G280 | 269.8 | 45 | -0.06 | 0.10 | 0.12 | 1 | 46 | 5.5 | 3 |
| G275 | 276.9 | 36 | 0.04 | -0.88 | 0.88 | 13 | 49 | 5.4 | 2 |
| G289 | 268.5 | 48 | -0.38 | -0.08 | 0.45 | 5 | 53 | 6.2 | 29 |
| G254 | 273.5 | 41 | -0.75 | -1.30 | 1.57 | 38 | 79 | 6.3 | 33 |
| G269 | 266.3 | 52 | 1.18 | -0.40 | 1.43 | 28 | 80 | 6.7 | 56 |
| G156 | 265 | 59 | -0.57 | 1.13 | 1.31 | 24 | 83 | 6.9 | 63 |
| G177 | 256.6 | 82 | 0.38 | 0.40 | 0.60 | 6 | 88 | 5.7 | 5 |
| G279 | 260.3 | 71 | 0.80 | 0.59 | 1.11 | 19 | 90 | 6.3 | 34 |
| G112 | 288.2 | 28 | -1.57 | -0.77 | 1.98 | 64 | 92 | 7.5 | 113 |
| G239 | 275.6 | 39 | 1.32 | 0.89 | 1.78 | 53 | 92 | 6.2 | 30 |
| G236 | 262.5 | 65 | -1.19 | -0.22 | 1.40 | 27 | 92 | 6.3 | 36 |
| G36 | 255.6 | 85 | 0.46 | -0.39 | 0.66 | 7 | 92 | 8.6 | 174 |
| G264 | 266.3 | 53 | 0.14 | 1.68 | 1.69 | 43 | 96 | 6.5 | 49 |
| G170 | 264.5 | 60 | -0.65 | 1.36 | 1.55 | 36 | 96 | 6.0 | 23 |
| G252 | 267.7 | 49 | -0.29 | 1.71 | 1.74 | 50 | 99 | 5.9 | 16 |
| G215 | 253.8 | 89 | 0.63 | 0.20 | 0.76 | 10 | 99 | 6.4 | 44 |
| G82 | 276.6 | 37 | 0.91 | -1.74 | 2.04 | 65 | 102 | 6.5 | 51 |
| G173 | 266 | 56 | -0.05 | 1.72 | 1.72 | 47 | 103 | 6.2 | 26 |
| G291 | 263.3 | 63 | -1.39 | 0.35 | 1.66 | 41 | 104 | 7.5 | 110 |
| G61 | 266.9 | 51 | 0.67 | 1.63 | 1.81 | 55 | 106 | 7.7 | 127 |
| G46 | 250.4 | 103 | 0.05 | 0.19 | 0.20 | 3 | 106 | 5.8 | 11 |
| G267 | 259.6 | 73 | 1.26 | -0.37 | 1.51 | 35 | 108 | 6.7 | 55 |
| G262 | 265.7 | 58 | -0.14 | -1.76 | 1.77 | 52 | 110 | 6.3 | 41 |
| G27 | 254.1 | 88 | -0.17 | 1.29 | 1.30 | 23 | 111 | 8.5 | 172 |
| G206 | 276.4 | 38 | -0.83 | 2.05 | 2.27 | 75 | 113 | 5.7 | 8 |
| G83 | 248.5 | 109 | 0.05 | 0.43 | 0.44 | 4 | 113 | 7.9 | 142 |
| G231 | 271.4 | 43 | -1.13 | 1.73 | 2.17 | 72 | 115 | 5.9 | 14 |
| G2 | 259 | 76 | 1.24 | -0.74 | 1.62 | 39 | 115 | 6.4 | 43 |
| G218 | 266.3 | 54 | 0.75 | 1.77 | 1.98 | 63 | 117 | 6.3 | 38 |
| **Least stable** | |  |  |  |  |  |  |  |  |
| G150 | 196 | 194 | -0.59 | -7.85 | 7.88 | 180 | 374 | 8.5 | 170 |
| G73 | 198.7 | 193 | 0.69 | -6.11 | 6.16 | 172 | 365 | 7.7 | 130 |
| G148 | 200.3 | 191 | 2.36 | -4.65 | 5.40 | 161 | 352 | 8.7 | 177 |
| G86 | 212.5 | 178 | 3.29 | -4.97 | 6.28 | 173 | 351 | 7.8 | 134 |
| G248 | 215.1 | 174 | 0.52 | -5.98 | 6.01 | 170 | 344 | 7.4 | 89 |
| G181 | 217.1 | 173 | 2.55 | -5.14 | 5.94 | 169 | 342 | 7.8 | 135 |
| G119 | 210 | 184 | 4.11 | 0.06 | 4.79 | 151 | 335 | 8.1 | 152 |
| G178 | 220 | 168 | 0.12 | -5.54 | 5.54 | 163 | 331 | 10.7 | 195 |
| G257 | 210.6 | 183 | 3.13 | -2.86 | 4.64 | 146 | 329 | 7.7 | 125 |
| G72 | 199 | 192 | 0.05 | -4.11 | 4.11 | 135 | 327 | 8.1 | 155 |
| G110 | 212.9 | 177 | 4.01 | 0.83 | 4.75 | 148 | 325 | 8.7 | 176 |
| G48 | 185.2 | 196 | 2.85 | -1.53 | 3.66 | 127 | 323 | 9.5 | 192 |
| G283 | 211.7 | 180 | 3.05 | -2.73 | 4.48 | 143 | 323 | 7.7 | 131 |
| G34 | 206.1 | 187 | 0.98 | -3.91 | 4.07 | 133 | 320 | 9.3 | 189 |
| G87 | 211.6 | 181 | 2.65 | -2.68 | 4.10 | 134 | 315 | 8.8 | 183 |
| G166 | 232.1 | 147 | 2.03 | -5.33 | 5.83 | 167 | 314 | 7.7 | 132 |
| G118 | 226.6 | 156 | 3.72 | -2.61 | 5.06 | 157 | 313 | 7.5 | 108 |
| G4 | 224.8 | 160 | 1.16 | 4.58 | 4.78 | 150 | 310 | 8.9 | 186 |
| G169 | 234.3 | 143 | 0.23 | -5.80 | 5.80 | 166 | 309 | 6.0 | 22 |
| G217 | 225.6 | 158 | 0.76 | -4.69 | 4.77 | 149 | 307 | 7.3 | 83 |
| G161 | 223.5 | 161 | 1.31 | -4.36 | 4.62 | 145 | 306 | 7.4 | 93 |
| G12 | 214.8 | 175 | 2.77 | -1.63 | 3.62 | 126 | 301 | 8.3 | 160 |
| G151 | 222.7 | 163 | -0.07 | -4.13 | 4.13 | 137 | 300 | 7.5 | 104 |
| G276 | 221.5 | 166 | 3.10 | -1.77 | 4.03 | 132 | 298 | 7.5 | 115 |
| G149 | 206.6 | 186 | 1.45 | -2.43 | 2.96 | 109 | 295 | 7.3 | 87 |
| G282 | 211.9 | 179 | 1.48 | -2.66 | 3.17 | 116 | 295 | 7.2 | 80 |
| G221 | 213.2 | 176 | 2.74 | 1.02 | 3.35 | 119 | 295 | 8.2 | 156 |
| G31 | 222.5 | 164 | 3.29 | 0.46 | 3.86 | 130 | 294 | 8.7 | 178 |
| G8 | 202.5 | 190 | 2.47 | 0.02 | 2.88 | 103 | 293 | 8.8 | 182 |
| G32 | 234.9 | 139 | 4.11 | 0.95 | 4.88 | 154 | 293 | 7.9 | 141 |

| Supplemental Table 5: p values for Likelihood Ratio Test of the analysed traits. | | | | | |
| --- | --- | --- | --- | --- | --- |
| **Traits** | **DH** | **DM** | **PH** | **TKW** | **GY** |
| **Genotype** | 8.95e-14** | 2.21e-14** | 4.38e-71** | 7.36e-55** | 2.65e-61** |
| **Genotype × Environment** | 4.50e-32** | 3.57e-34** | 1.38e-14** | 1.26e-115** | 1.58e-93** |
| **variables with p<0.01 represent highly significant. | | | | | |

| Supplemental Table 6: Eigenvalues, explained variance, factorial loadings after varimax rotation, and communalities and uniqueness obtained in the factor analysis of the 5 variables studied in 196 wheat genotypes across twelve environments during *Rabi* 2020-21 and 2021-22. | | | | | | |
| --- | --- | --- | --- | --- | --- | --- |
|  | **VAR** | **FA1** | **FA2** | **FA3** | **Communality** | **Uniqueness** |
| **1** | **DH** | **-0.85** | -0.13 | 0.10 | 0.74 | 0.26 |
| **2** | **DM** | **-0.87** | 0.005 | -0.05 | 0.76 | 0.23 |
| **3** | **PH** | -0.071 | **0.80** | 0.20 | 0.69 | 0.30 |
| **4** | **TKW** | -0.20 | **0.73** | -0.20 | 0.62 | 0.37 |
| **5** | **GY** | -0.022 | 0.012 | **0.97** | 0.93 | 0.06 |
|  | **Eigenvalues** | 1.63 | 1.14 | 0.98 |  |  |
|  | **Variance %** | 32.67 | 22.85 | 19.7 |  |  |
|  | **Cumulative variance %** | 32.67 | 55.52 | 75.22 |  |  |
| DH, days to heading; DM, days to maturity; PH, plant height; TKW, thousand kernel weight; GY, grain yield.  Bold values indicated the variables grouped within each factor FA. | | | | | | |

| Supplemental Table 7: Genotype-ideotype (ID) score, MTSI values for the 196 wheat genotypes for the three factors along with relative contribution of each factor towards the MTSI evaluated at twelve environments during 2020-21 and 2021-22. | | | | | | | | | | | | | | | |
| --- | --- | --- | --- | --- | --- | --- | --- | --- | --- | --- | --- | --- | --- | --- | --- |
| **Entry No** | **$FA1** | | **FA2** | | **FA3** | | **MTSI** | **Entry No** | **FA1** | | **FA2** | | **FA3** | | **MTSI** |
|  | **Score** | **RC%** | **Score** | **RC%** | **Score** | **RC%** |  |  | **Score** | **RC%** | **Score** | **RC%** | **Score** | **RC%** |  |
| G2 | -6.6 | 47.8 | 8.6 | 36.1 | 4.6 | 16.1 | 6.2 | G161 | -9.2 | 20.7 | 7.1 | 46.8 | 2.7 | 32.4 | 6.6 |
| G4 | -7.1 | 32.7 | 6.3 | 44.0 | 3.1 | 23.3 | 8.0 | G165 | -8.0 | 31.4 | 7.4 | 43.8 | 3.5 | 24.7 | 6.6 |
| G8 | -7.2 | 29.1 | 6.9 | 36.3 | 1.2 | 34.6 | 8.5 | G166 | -7.7 | 34.6 | 8.5 | 35.2 | 3.0 | 30.1 | 6.2 |
| G9 | -7.5 | 30.1 | 6.2 | 46.1 | 3.1 | 23.8 | 7.8 | G168 | -7.9 | 35.0 | 8.3 | 38.8 | 3.6 | 26.2 | 5.9 |
| G12 | -7.3 | 36.6 | 8.2 | 35.9 | 3.1 | 27.6 | 6.6 | **G169** | **-9.3** | **26.3** | **8.9** | **40.2** | **3.4** | **33.5** | **4.8** |
| G16 | -7.3 | 31.9 | 6.9 | 40.9 | 2.7 | 27.3 | 7.6 | G170 | -7.8 | 34.9 | 6.6 | 53.7 | 5.0 | 11.4 | 6.8 |
| G19 | -7.3 | 36.8 | 8.1 | 36.5 | 3.2 | 26.7 | 6.6 | **G172** | **-8.3** | **35.2** | **8.1** | **46.7** | **4.6** | **18.1** | **5.4** |
| G20 | -9.2 | 23.5 | 8.1 | 43.4 | 3.0 | 33.1 | 5.7 | **G173** | **-8.6** | **35.2** | **9.1** | **38.6** | **4.1** | **26.2** | **4.7** |
| **G21** | **-8.7** | **33.3** | **8.5** | **45.5** | **4.5** | **21.2** | **4.9** | G174 | -8.6 | 32.0 | 7.2 | 57.2 | 5.3 | 10.8 | 5.8 |
| G23 | -7.8 | 28.7 | 5.3 | 54.3 | 4.0 | 17.0 | 8.2 | **G175** | **-8.7** | **39.1** | **8.1** | **59.5** | **6.1** | **1.4** | **4.9** |
| G25 | -7.3 | 41.7 | 8.3 | 39.5 | 4.3 | 18.8 | 6.0 | G177 | -7.7 | 37.7 | 8.9 | 33.1 | 3.3 | 29.2 | 5.8 |
| G26 | -7.4 | 29.8 | 6.2 | 44.6 | 2.8 | 25.6 | 8.0 | G178 | -5.3 | 48.0 | 7.8 | 34.9 | 4.0 | 17.0 | 7.9 |
| G27 | -6.4 | 37.0 | 6.7 | 40.8 | 3.2 | 22.1 | 8.0 | G179 | -7.3 | 46.5 | 8.1 | 45.7 | 5.5 | 7.8 | 5.9 |
| G29 | -7.9 | 28.4 | 6.4 | 46.6 | 3.1 | 25.0 | 7.5 | G181 | -6.5 | 45.2 | 8.6 | 33.9 | 3.9 | 21.0 | 6.5 |
| G30 | -6.9 | 32.1 | 6.7 | 39.1 | 2.1 | 28.8 | 8.3 | G183 | -7.7 | 38.4 | 7.8 | 45.9 | 4.7 | 15.7 | 6.0 |
| G31 | -8.0 | 26.7 | 6.3 | 46.7 | 2.8 | 26.5 | 7.7 | G186 | -6.3 | 42.6 | 7.4 | 40.5 | 4.2 | 16.9 | 7.3 |
| G32 | -8.8 | 24.8 | 7.7 | 42.3 | 2.7 | 32.8 | 6.3 | **G187** | **-8.4** | **37.4** | **8.0** | **52.1** | **5.4** | **10.5** | **5.3** |
| G34 | -7.4 | 32.7 | 7.1 | 41.2 | 3.0 | 26.1 | 7.3 | **G188** | **-8.2** | **36.3** | **8.6** | **40.9** | **4.2** | **22.7** | **5.3** |
| G35 | -8.4 | 23.5 | 5.7 | 50.2 | 2.8 | 26.3 | 8.0 | G189 | -8.3 | 29.5 | 7.1 | 48.2 | 3.8 | 22.3 | 6.5 |
| G36 | -8.6 | 23.5 | 5.5 | 55.1 | 3.6 | 21.4 | 7.8 | G190 | -8.2 | 28.7 | 5.9 | 56.5 | 4.5 | 14.9 | 7.3 |
| G37 | -7.0 | 31.8 | 6.4 | 42.5 | 2.7 | 25.7 | 8.1 | **G191** | **-9.1** | **25.4** | **8.3** | **43.5** | **3.4** | **31.1** | **5.4** |
| G39 | -7.6 | 34.9 | 7.3 | 44.8 | 4.0 | 20.3 | 6.6 | **G192** | **-10.2** | **16.9** | **7.9** | **58.3** | **4.4** | **24.8** | **4.9** |
| G40 | -8.1 | 34.0 | 8.1 | 41.6 | 3.8 | 24.4 | 5.8 | G195 | -8.5 | 31.6 | 7.7 | 49.7 | 4.5 | 18.7 | 5.6 |
| G43 | -6.6 | 42.7 | 7.8 | 39.0 | 4.2 | 18.2 | 6.8 | **G198** | **-8.8** | **29.0** | **8.1** | **46.1** | **4.0** | **24.8** | **5.3** |
| G44 | -9.0 | 27.4 | 7.1 | 57.5 | 4.9 | 15.1 | 5.8 | G200 | -8.8 | 28.1 | 8.1 | 44.8 | 3.7 | 27.2 | 5.5 |
| **G45** | **-11.0** | **6.0** | **7.6** | **63.1** | **3.9** | **30.9** | **5.2** | G201 | -8.4 | 29.0 | 7.3 | 47.8 | 3.8 | 23.2 | 6.3 |
| G46 | -8.3 | 31.7 | 7.7 | 45.0 | 3.9 | 23.4 | 6.0 | G202 | -7.4 | 39.5 | 7.2 | 49.6 | 5.1 | 10.9 | 6.6 |
| G47 | -7.6 | 31.9 | 6.3 | 49.0 | 3.9 | 19.2 | 7.4 | G203 | -8.5 | 29.0 | 6.2 | 59.6 | 5.1 | 11.5 | 6.8 |
| G48 | -7.6 | 31.6 | 7.0 | 43.0 | 3.1 | 25.4 | 7.2 | G204 | -8.4 | 28.2 | 6.8 | 51.3 | 4.0 | 20.5 | 6.6 |
| G49 | -8.0 | 33.3 | 6.5 | 56.1 | 5.1 | 10.6 | 6.8 | G205 | -9.4 | 23.9 | 6.7 | 64.7 | 5.2 | 11.3 | 6.0 |
| G54 | -8.8 | 26.5 | 5.8 | 64.7 | 5.3 | 8.8 | 7.0 | G206 | -9.9 | 18.3 | 7.1 | 60.0 | 4.4 | 21.7 | 5.7 |
| G56 | -8.9 | 28.6 | 6.8 | 60.3 | 5.2 | 11.1 | 6.1 | G207 | -6.5 | 38.6 | 6.9 | 41.9 | 3.7 | 19.5 | 7.7 |
| G57 | -7.2 | 40.6 | 6.7 | 53.2 | 5.6 | 6.2 | 7.0 | **G209** | **-8.3** | **37.1** | **9.1** | **37.1** | **4.0** | **25.8** | **5.0** |
| G58 | -7.5 | 33.1 | 6.9 | 44.8 | 3.6 | 22.1 | 7.2 | G210 | -8.2 | 32.6 | 8.8 | 34.4 | 3.0 | 32.9 | 5.7 |
| G59 | -6.9 | 37.6 | 7.2 | 41.8 | 3.7 | 20.7 | 7.3 | G211 | -6.5 | 42.9 | 9.2 | 27.0 | 2.8 | 30.1 | 6.7 |
| G61 | -7.3 | 33.2 | 6.8 | 42.8 | 3.2 | 24.0 | 7.4 | **G212** | **-7.9** | **39.7** | **8.7** | **39.7** | **4.4** | **20.7** | **5.4** |
| G62 | -7.4 | 36.6 | 7.8 | 40.6 | 3.7 | 22.7 | 6.5 | **G213** | **-10.5** | **13.9** | **7.5** | **71.8** | **5.3** | **14.3** | **4.9** |
| G63 | -6.9 | 39.1 | 7.0 | 44.5 | 4.3 | 16.4 | 7.2 | G215 | -8.5 | 29.6 | 8.0 | 41.9 | 3.4 | 28.4 | 5.8 |
| G64 | -6.8 | 37.0 | 7.8 | 35.5 | 2.8 | 27.5 | 7.2 | G216 | -7.6 | 41.5 | 8.1 | 44.4 | 4.9 | 14.1 | 5.8 |
| G65 | -8.1 | 28.4 | 7.0 | 45.3 | 3.2 | 26.2 | 6.9 | G217 | -8.9 | 24.3 | 7.5 | 45.6 | 3.1 | 30.0 | 6.2 |
| G66 | -7.5 | 35.7 | 7.6 | 41.8 | 3.7 | 22.4 | 6.6 | G218 | -8.1 | 36.4 | 8.6 | 38.7 | 3.9 | 24.9 | 5.4 |
| G67 | -7.6 | 37.0 | 7.5 | 45.6 | 4.4 | 17.4 | 6.4 | G219 | -7.3 | 43.1 | 8.0 | 43.7 | 4.9 | 13.2 | 6.0 |
| G68 | -7.1 | 34.1 | 7.7 | 35.3 | 2.3 | 30.7 | 7.4 | G220 | -7.9 | 33.0 | 7.0 | 49.2 | 4.3 | 17.8 | 6.6 |
| G69 | -6.5 | 36.3 | 6.7 | 40.6 | 3.0 | 23.1 | 8.1 | G221 | -9.6 | 18.2 | 7.0 | 50.5 | 3.0 | 31.3 | 6.4 |
| G71 | -8.4 | 24.4 | 7.4 | 39.2 | 1.8 | 36.4 | 7.2 | G222 | -8.9 | 23.4 | 7.1 | 47.7 | 3.1 | 28.9 | 6.5 |
| G72 | -9.4 | 17.5 | 7.5 | 40.0 | 1.2 | 42.6 | 7.2 | G225 | -6.4 | 39.2 | 8.9 | 26.1 | 1.8 | 34.8 | 7.5 |
| G73 | -9.2 | 22.0 | 7.0 | 51.6 | 3.6 | 26.4 | 6.2 | G226 | -7.4 | 38.5 | 8.1 | 39.2 | 3.9 | 22.3 | 6.2 |
| G74 | -5.1 | 42.6 | 5.4 | 45.9 | 4.5 | 11.5 | 9.5 | G227 | -9.0 | 24.5 | 7.6 | 46.8 | 3.3 | 28.7 | 6.0 |
| G75 | -8.7 | 21.3 | 6.8 | 43.2 | 1.7 | 35.6 | 7.5 | **G228** | **-8.8** | **30.4** | **8.3** | **45.8** | **4.2** | **23.9** | **5.1** |
| G76 | -7.7 | 30.2 | 6.6 | 45.1 | 3.1 | 24.8 | 7.4 | G231 | -3.5 | 59.7 | 9.1 | 23.7 | 4.0 | 16.6 | 8.8 |
| G78 | -8.5 | 26.7 | 6.8 | 49.1 | 3.5 | 24.3 | 6.7 | **G232** | **-9.2** | **24.4** | **8.5** | **40.6** | **3.0** | **35.0** | **5.4** |
| G80 | -8.2 | 33.6 | 7.8 | 45.8 | 4.2 | 20.7 | 5.8 | G235 | -9.0 | 25.9 | 9.2 | 32.0 | 2.2 | 42.1 | 5.6 |
| G82 | -9.2 | 19.5 | 7.0 | 44.7 | 2.1 | 35.8 | 7.0 | G236 | -8.0 | 32.9 | 8.1 | 40.3 | 3.5 | 26.8 | 6.0 |
| G83 | -7.9 | 28.6 | 5.6 | 52.9 | 3.9 | 18.5 | 7.8 | G237 | -10.8 | 7.3 | 6.2 | 65.6 | 3.7 | 27.1 | 6.6 |
| **G84** | **-7.9** | **41.4** | **9.0** | **38.3** | **4.5** | **20.3** | **5.1** | G239 | -7.3 | 34.9 | 7.5 | 40.5 | 3.3 | 24.6 | 6.9 |
| **G86** | **-8.9** | **28.5** | **9.1** | **36.1** | **3.1** | **35.4** | **5.1** | G240 | -7.6 | 34.5 | 7.1 | 46.0 | 4.0 | 19.6 | 6.8 |
| G87 | -8.7 | 25.5 | 7.3 | 45.5 | 3.1 | 28.9 | 6.5 | **G241** | **-8.1** | **37.4** | **8.8** | **38.0** | **4.0** | **24.6** | **5.3** |
| G88 | -8.4 | 21.8 | 4.6 | 55.1 | 3.0 | 23.1 | 8.8 | G244 | -9.3 | 20.7 | 7.3 | 48.4 | 3.0 | 30.9 | 6.3 |
| G89 | -8.6 | 22.6 | 5.2 | 55.8 | 3.5 | 21.5 | 8.1 | G248 | -7.2 | 38.1 | 8.4 | 34.6 | 3.2 | 27.3 | 6.5 |
| G91 | -7.7 | 27.9 | 7.4 | 35.9 | 1.4 | 36.2 | 7.8 | **G249** | **-9.4** | **28.3** | **10.8** | **20.2** | **2.5** | **51.5** | **4.5** |
| G92 | -8.5 | 24.3 | 5.6 | 54.4 | 3.6 | 21.3 | 7.7 | G250 | -8.6 | 22.7 | 7.8 | 35.5 | 1.0 | 41.8 | 7.5 |
| G94 | -8.4 | 29.2 | 7.4 | 45.9 | 3.6 | 24.9 | 6.3 | G251 | -7.2 | 32.9 | 8.2 | 31.2 | 1.6 | 35.9 | 7.4 |
| G95 | -8.4 | 28.3 | 7.2 | 46.7 | 3.5 | 25.1 | 6.5 | G252 | -8.6 | 25.4 | 8.7 | 31.8 | 1.5 | 42.9 | 6.5 |
| G96 | -8.5 | 21.7 | 6.3 | 44.5 | 1.7 | 33.8 | 8.0 | G253 | -6.8 | 41.6 | 8.4 | 33.8 | 3.5 | 24.5 | 6.6 |
| G99 | -9.4 | 22.1 | 6.8 | 58.2 | 4.4 | 19.7 | 6.1 | G254 | -7.2 | 36.2 | 6.5 | 49.2 | 4.5 | 14.6 | 7.3 |
| G101 | -8.5 | 23.5 | 7.1 | 41.6 | 1.9 | 34.9 | 7.3 | G256 | -7.1 | 40.4 | 8.3 | 36.7 | 3.8 | 22.9 | 6.3 |
| G102 | -8.6 | 23.9 | 8.1 | 34.9 | 1.3 | 41.2 | 7.0 | G257 | -7.9 | 32.0 | 8.2 | 36.6 | 2.8 | 31.4 | 6.3 |
| **G103** | **-9.3** | **29.4** | **7.5** | **64.4** | **5.8** | **6.2** | **5.2** | G258 | -7.3 | 32.6 | 7.3 | 39.2 | 2.6 | 28.2 | 7.4 |
| G105 | -8.3 | 22.7 | 6.3 | 43.3 | 1.6 | 34.0 | 8.1 | G259 | -7.1 | 35.9 | 6.6 | 46.6 | 4.1 | 17.6 | 7.4 |
| G106 | -8.2 | 31.7 | 8.0 | 41.5 | 3.5 | 26.7 | 6.0 | G261 | -7.3 | 38.7 | 8.0 | 39.7 | 3.9 | 21.6 | 6.4 |
| G107 | -8.1 | 27.3 | 5.9 | 51.3 | 3.6 | 21.4 | 7.7 | G262 | -8.7 | 30.1 | 7.9 | 47.8 | 4.2 | 22.1 | 5.5 |
| G108 | -7.7 | 24.5 | 4.0 | 54.0 | 2.9 | 21.5 | 9.6 | G264 | -6.7 | 41.6 | 7.7 | 39.6 | 4.1 | 18.8 | 6.9 |
| G110 | -8.0 | 27.9 | 5.9 | 51.5 | 3.7 | 20.6 | 7.7 | G265 | -8.3 | 34.1 | 7.9 | 46.1 | 4.4 | 19.8 | 5.6 |
| G112 | -5.9 | 42.6 | 6.8 | 41.8 | 4.2 | 15.6 | 8.0 | G266 | -8.2 | 33.2 | 7.8 | 44.8 | 4.1 | 22.0 | 5.9 |
| **G113** | **-8.6** | **30.6** | **8.8** | **36.9** | **3.2** | **32.5** | **5.4** | **G267** | **-8.2** | **36.9** | **9.2** | **34.1** | **3.7** | **28.9** | **5.1** |
| G114 | -9.2 | 21.1 | 6.0 | 58.8 | 4.1 | 20.1 | 6.9 | G268 | -8.2 | 33.2 | 7.3 | 50.9 | 4.7 | 15.9 | 6.0 |
| G115 | -7.9 | 26.8 | 7.3 | 37.2 | 1.5 | 35.9 | 7.7 | G269 | -6.8 | 43.7 | 8.3 | 37.1 | 4.2 | 19.2 | 6.3 |
| G116 | -7.8 | 34.6 | 7.3 | 46.4 | 4.2 | 19.0 | 6.5 | G272 | -6.2 | 50.4 | 7.3 | 47.7 | 6.4 | 1.9 | 7.2 |
| G117 | -8.0 | 32.4 | 8.1 | 38.0 | 3.0 | 29.5 | 6.2 | G273 | -6.1 | 45.7 | 8.1 | 35.2 | 4.0 | 19.1 | 7.1 |
| G118 | -7.9 | 36.9 | 8.5 | 39.1 | 3.9 | 24.0 | 5.6 | G274 | -8.6 | 35.4 | 7.6 | 58.0 | 5.7 | 6.6 | 5.5 |
| G119 | -8.4 | 30.0 | 8.4 | 37.6 | 2.9 | 32.4 | 5.9 | **G275** | **-8.0** | **42.0** | **8.8** | **42.3** | **4.9** | **15.7** | **5.0** |
| G123 | -7.9 | 27.4 | 7.4 | 37.3 | 1.6 | 35.2 | 7.6 | **G276** | **-9.4** | **23.7** | **8.5** | **42.7** | **3.3** | **33.6** | **5.2** |
| **G126** | **-8.7** | **29.9** | **8.3** | **42.6** | **3.7** | **27.5** | **5.4** | G277 | -7.1 | 39.2 | 7.8 | 39.9 | 3.9 | 20.9 | 6.6 |
| G128 | -7.5 | 33.2 | 7.2 | 42.1 | 3.3 | 24.7 | 7.0 | G278 | -6.1 | 46.1 | 8.4 | 33.0 | 3.8 | 20.9 | 7.0 |
| G129 | -6.9 | 32.1 | 6.9 | 37.7 | 2.0 | 30.2 | 8.2 | G279 | -6.4 | 42.0 | 7.3 | 41.7 | 4.3 | 16.3 | 7.3 |
| G132 | -7.6 | 37.9 | 8.5 | 37.0 | 3.7 | 25.1 | 5.9 | G280 | -9.0 | 25.3 | 7.7 | 46.5 | 3.5 | 28.2 | 5.9 |
| G133 | -7.2 | 33.3 | 6.6 | 44.4 | 3.4 | 22.3 | 7.7 | **G281** | **-9.2** | **27.5** | **9.3** | **36.3** | **3.3** | **36.2** | **4.7** |
| G135 | -8.5 | 25.4 | 5.9 | 54.8 | 3.9 | 19.8 | 7.3 | G282 | -8.0 | 28.0 | 7.2 | 41.5 | 2.5 | 30.5 | 7.1 |
| G136 | -8.0 | 26.0 | 5.1 | 53.0 | 3.4 | 21.0 | 8.4 | G283 | -8.1 | 33.2 | 8.4 | 38.1 | 3.3 | 28.7 | 5.9 |
| G138 | -8.0 | 31.4 | 7.0 | 47.4 | 3.9 | 21.3 | 6.7 | G285 | -7.6 | 32.1 | 7.8 | 37.2 | 2.5 | 30.7 | 7.0 |
| G141 | -8.7 | 27.0 | 7.2 | 49.6 | 3.9 | 23.3 | 6.2 | **G286** | **-8.0** | **39.3** | **8.9** | **38.2** | **4.2** | **22.4** | **5.2** |
| G148 | -8.5 | 25.1 | 6.6 | 48.1 | 3.1 | 26.8 | 7.1 | G289 | -7.2 | 41.3 | 8.1 | 40.3 | 4.3 | 18.4 | 6.2 |
| G149 | -9.6 | 17.8 | 6.9 | 53.1 | 3.3 | 29.1 | 6.4 | G290 | -7.9 | 33.5 | 7.6 | 44.6 | 3.9 | 21.9 | 6.3 |
| G150 | -8.7 | 28.5 | 8.5 | 38.4 | 3.0 | 33.1 | 5.7 | G291 | -6.9 | 34.9 | 6.6 | 43.2 | 3.4 | 21.8 | 7.8 |
| G151 | -7.5 | 34.2 | 7.6 | 40.2 | 3.2 | 25.6 | 6.8 | G292 | -8.5 | 30.9 | 8.3 | 41.8 | 3.6 | 27.3 | 5.6 |
| G155 | -8.9 | 22.5 | 6.2 | 53.0 | 3.4 | 24.5 | 7.2 | G293 | -7.2 | 30.8 | 5.7 | 46.9 | 3.1 | 22.4 | 8.4 |
| G156 | -8.4 | 30.6 | 7.2 | 50.1 | 4.3 | 19.3 | 6.1 | G294 | -6.5 | 34.2 | 7.9 | 30.0 | 1.0 | 35.8 | 8.4 |
| G157 | -8.7 | 26.1 | 7.4 | 45.9 | 3.2 | 28.0 | 6.3 | **ID** | **11.3** |  | **-10.8** |  | **11.0** |  | **0.0** |
| **G158** | **-8.9** | **30.5** | **7.9** | **52.5** | **4.8** | **17.0** | **5.2** |  |  |  |  |  |  |  |  |

RC (%), relative contribution; MTSI, multi-trait stability index; ID, ideotype;

$Factors retained

Bold values indicate selected genotypes with lower MTSI values.

| Supplemental Table 8: Mean grain yield, WAASB values of 63 genotypes grouped under fourth quadrant of WAASBY bi-plot (Fig. 6), representing most stable and having mean grain yield higher than grand mean of the population at twelve environments during *Rabi* 2020-21 and 2021-22. | | | | | | | | | |
| --- | --- | --- | --- | --- | --- | --- | --- | --- | --- |
| **Sl. No.** | **Entry** | **Grain yield Mean** | **WAASB** | **QUADRANT** | **Sl. No.** | **Entry** | **Grain yield Mean** | **WAASB** | **QUADRANT** |
| 1 | G175 | 488.2 | 1.5 | 4 | 33 | G289 | 364.8 | 0.8 | 4 |
| 2 | G103 | 473.3 | 1.4 | 4 | 34 | G198 | 364.8 | 1.3 | 4 |
| 3 | G272 | 471.0 | 0.9 | 4 | 35 | G39 | 363.0 | 1.2 | 4 |
| 4 | G54 | 452.3 | 1.3 | 4 | 36 | G80 | 360.0 | 0.8 | 4 |
| 5 | G57 | 451.1 | 1.0 | 4 | 37 | G218 | 359.9 | 1.2 | 4 |
| 6 | G187 | 448.8 | 1.4 | 4 | 38 | G212 | 359.8 | 1.0 | 4 |
| 7 | G174 | 437.3 | 1.5 | 4 | 39 | G158 | 359.1 | 0.5 | 4 |
| 8 | G274 | 432.1 | 1.2 | 4 | 40 | G264 | 357.1 | 1.5 | 4 |
| 9 | G216 | 431.5 | 1.3 | 4 | 41 | G107 | 356.9 | 1.5 | 4 |
| 10 | G56 | 430.2 | 0.9 | 4 | 42 | G2 | 356.7 | 0.8 | 4 |
| 11 | G179 | 428.8 | 1.0 | 4 | 43 | G183 | 356.7 | 0.7 | 4 |
| 12 | G205 | 413.2 | 0.8 | 4 | 44 | G170 | 356.2 | 0.5 | 4 |
| 13 | G219 | 412.5 | 0.7 | 4 | 45 | G269 | 353.1 | 1.3 | 4 |
| 14 | G44 | 408.6 | 0.8 | 4 | 46 | G190 | 352.9 | 1.2 | 4 |
| 15 | G213 | 408.5 | 0.4 | 4 | 47 | G192 | 352.8 | 0.7 | 4 |
| 16 | G25 | 407.0 | 1.3 | 4 | 48 | G279 | 352.1 | 0.7 | 4 |
| 17 | G99 | 401.9 | 1.2 | 4 | 49 | G278 | 351.7 | 1.3 | 4 |
| 18 | G63 | 395.9 | 1.2 | 4 | 50 | G292 | 351.5 | 1.5 | 4 |
| 19 | G112 | 388.7 | 1.4 | 4 | 51 | G67 | 351.2 | 0.7 | 4 |
| 20 | G228 | 382.9 | 1.2 | 4 | 52 | G231 | 351.1 | 0.9 | 4 |
| 21 | G268 | 381.3 | 1.3 | 4 | 53 | G108 | 350.6 | 1.1 | 4 |
| 22 | G275 | 380.1 | 1.0 | 4 | 54 | G62 | 350.0 | 1.3 | 4 |
| 23 | G47 | 379.5 | 1.5 | 4 | 55 | G156 | 349.2 | 1.0 | 4 |
| 24 | G21 | 377.0 | 1.2 | 4 | 56 | G241 | 348.3 | 0.8 | 4 |
| 25 | G84 | 373.9 | 1.1 | 4 | 57 | G173 | 348.1 | 0.9 | 4 |
| 26 | G138 | 372.5 | 1.1 | 4 | 58 | G204 | 347.8 | 1.3 | 4 |
| 27 | G265 | 371.8 | 0.9 | 4 | 59 | G36 | 347.7 | 1.1 | 4 |
| 28 | G188 | 370.2 | 1.2 | 4 | 60 | G116 | 347.4 | 0.8 | 4 |
| 29 | G206 | 369.9 | 0.9 | 4 | 61 | G132 | 345.5 | 0.9 | 4 |
| 30 | G254 | 369.7 | 1.1 | 4 | 62 | G135 | 345.1 | 0.7 | 4 |
| 31 | G220 | 365.8 | 0.9 | 4 | 63 | G277 | 344.4 | 0.9 | 4 |
| 32 | G262 | 365.7 | 1.2 | 4 |  |  |  |  |  |

Supplemental Table 9: Mean, Standard deviation and standard error obtained under BLUP and AMMI family models using RMPD estimates.

| **MODEL** | **Mean** | **SD** | **SE** | **Q2.5** | **Q97.5** |
| --- | --- | --- | --- | --- | --- |
| BLUP | 70.5 | 5.83 | 0.825 | 62.3 | 83.4 |
| AMMIF | 71.7 | 8.42 | 1.19 | 55 | 85.8 |
| AMMI10 | 72 | 7.84 | 1.11 | 58.2 | 84.3 |
| AMMI7 | 72.1 | 7.64 | 1.08 | 57.3 | 85.3 |
| AMMI9 | 72.2 | 7.23 | 1.02 | 58.3 | 84.6 |
| AMMI8 | 72.2 | 6.04 | 0.855 | 59.8 | 81.9 |
| AMMI6 | 72.4 | 7.25 | 1.03 | 60.4 | 85.2 |
| AMMI3 | 73.3 | 3.52 | 0.497 | 68.1 | 81.6 |
| AMMI4 | 73.9 | 4.73 | 0.669 | 63.8 | 81.7 |
| AMMI5 | 73.9 | 6.83 | 0.966 | 60.8 | 85.7 |
| AMMI2 | 80.2 | 3.02 | 0.426 | 75.7 | 85.5 |
| AMMI1 | 91 | 2.47 | 0.349 | 88.3 | 97.3 |
| AMMI0 | 109 | 0.887 | 0.125 | 108 | 111 |

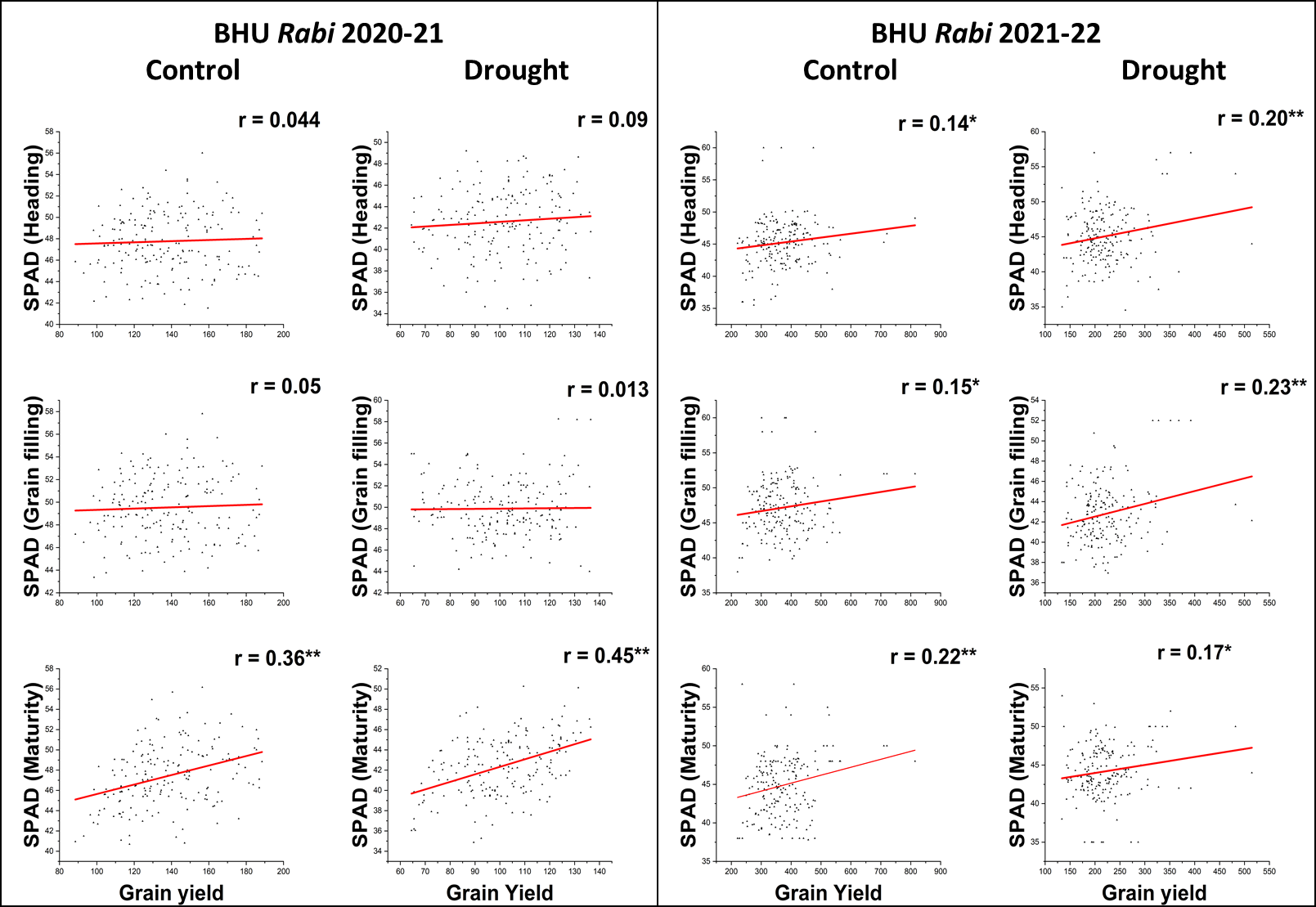

Supplementary Fig. 1. Regression analysis between grain yield vs SPAD readings under irrigated control and drought treatment evaluated at BHU location during *Rabi* 2020-21 and 2021-22. Observations were recorded during heading, grain filling and maturity. * and ** represent significant at 5% and 1% probability levels respectively.

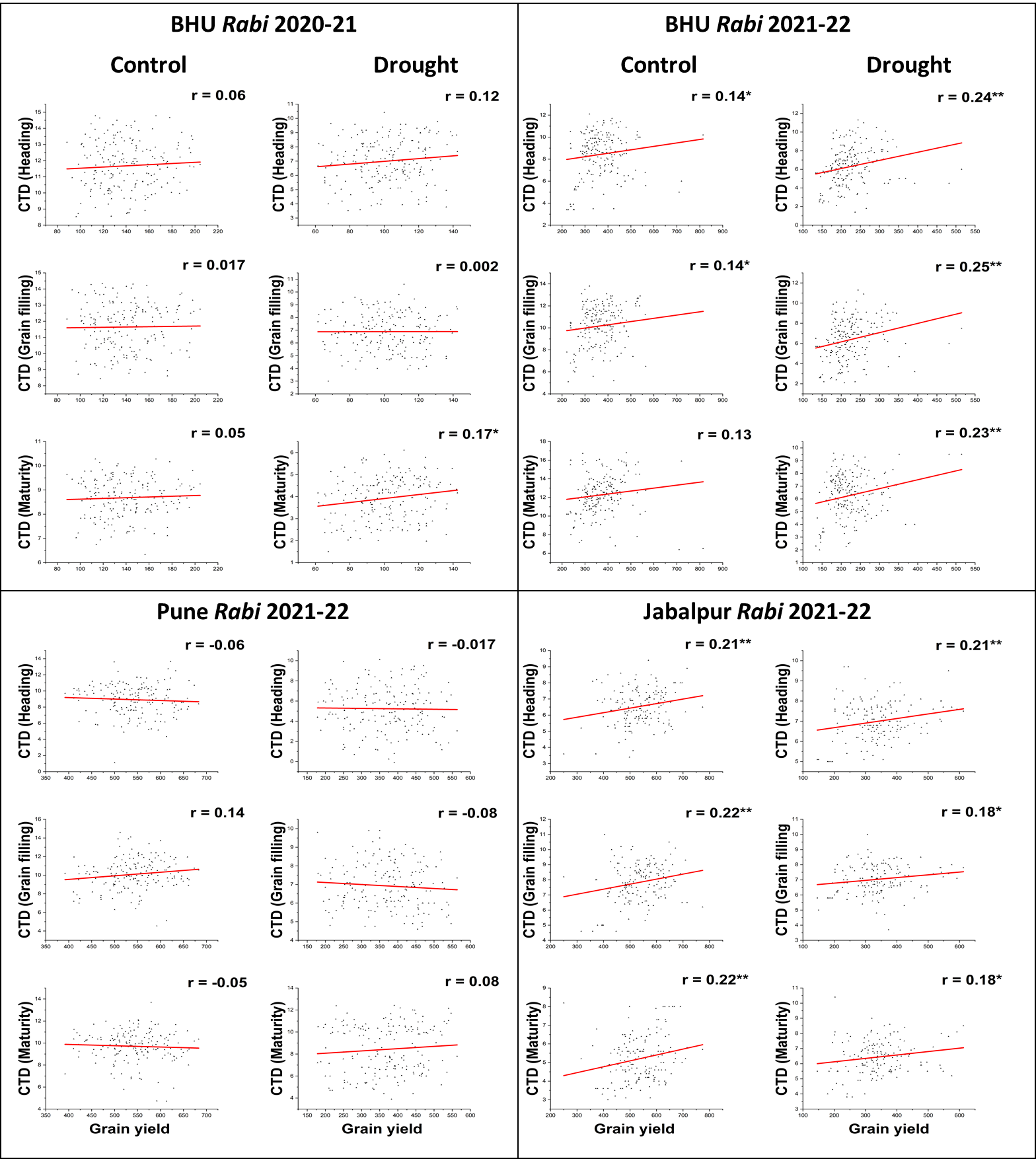

Supplementary Fig. 2. Regression analysis between grain yield vs CTD under irrigated control and drought at BHU during 2020-21 and 2021-22, Pune 2021-22 and Jabalpur 2021-22. CTD values recorded during heading, grain filling and maturity. * and ** represent significant at 5% and 1% probability levels respectively.
